# Forest decline differentially affects trophic guilds of canopy-dwelling beetles

**DOI:** 10.1101/2020.02.11.943753

**Authors:** Aurélien Sallé, Guilhem Parmain, Benoît Nusillard, Xavier Pineau, Ravène Brousse, Tiphanie Fontaine-Guenel, Romain Ledet, Cécile Vincent-Barbaroux, Christophe Bouget

## Abstract

**Context:** Decline in a context of climate change is expected to induce considerable changes in forest structure, potentially affecting habitat opportunities and trophic resources for numerous species. Nonetheless, the consequences of decline on forest biodiversity have rarely been studied.

**Aim:** We aimed to characterize the impact of oak decline on different guilds of canopy-dwelling beetles.

**Methods:** Beetles were sampled for three consecutive years in oak stands exhibiting different levels of decline. Several guilds were considered: (i) Buprestidae, (ii) other saproxylic beetles split into wood-boring species and non-wood-boring species, (iii) seed-eating weevils, and (iv) specialist and generalist leaf-eating weevils.

**Results:** Overall, decline had positive effects on the abundance and biomass of beetles, though contrasting variations were observed at the species or guild levels. Xylophagous species, especially the main oak-associated buprestids, and other saproxylic species benefitted from decline conditions. However, at odds with the insect performance hypothesis, decline had a positive effect on generalist phyllophagous species, a negative effect on specialist phyllophagous species, and a null effect on seminiphagous species.

**Conclusion:** The increase in species richness for saproxylic and phyllophagous beetle communities suggests that decline might promote forest biodiversity. Our results call for further studies to thoroughly assess the functional outcomes of forest decline, and to suggest management strategies for conservation biologists.

**Key message:** Decline can affect the structure, resources and microclimates of the forest canopy, and potentially have cascading effects on canopy-dwelling species. Our survey shows that an oak decline can promote saproxylic beetles, especially xylophagous ones, and generalist phyllophagous weevils. However, it negatively affects specialist phyllophagous species and has no effect on seminiphagous weevils.

## Introduction

Global change can dramatically affect the organization and functioning of forest ecosystems by promoting the introduction and establishment of invasive species (Liebhold et al. 2017), by intensifying land-use at the landscape level (Seibold et al. 2019), and through the direct and indirect effects of climate change on forest health (Seidl et al. 2017). Climate change already challenges the ability of European forests to adapt (Allen et al. 2010; Carnicer et al. 2011), and unprecedented forest declines are expected in response to the predicted increase in frequency and severity of droughts and heat waves (Allen et al. 2010; IPCC 2013).

Forest decline generally consists in a progressive loss of vigor of the trees, over several years, in response to multiple, successive or concomitant driving factors (Manion 1981). These factors include i) predisposing factors such as site conditions that constantly affect the stands, ii) inciting factors such as defoliation or droughts that trigger declines, and iii) contributing factors such as secondary pests and pathogens, which aggravate the deleterious effects of inciting factors, ultimately killing trees (Sinclair 1967; Manion 1981; Thomas et al. 2002; Sallé et al. 2014). The gradual loss of tree vigor progressively affects all forest compartments but the canopy is certainly the first to exhibit conspicuous modifications as decline progresses. The crown of a declining tree is characterized by an accumulation of dead branches, cavities and fruiting bodies of saprotrophic or pathogenic fungi (Houston 1981; Ishii et al. 2004). Therefore, a forest decline generates novel structures and favors the accumulation uncommon ones for healthy trees, and consequently tends to increase the structural complexity of the canopy at stand, tree and branch scales (Ishii et al. 2004). Crowns of declining trees also exhibit reduced foliage density, which in turn can considerably alter microclimates within and beneath the canopy (Houston 1981; Ishii et al. 2004). Such profound structural modifications affect habitat opportunities and trophic resources, with likely marked cascading effects on canopy-dwelling communities.

The tree canopy and the soil are the two key compartments supporting forest biodiversity and their contribution is tremendous (Stork and Grimbacher 2006). Compared to tropical forests, temperate forests have less vertical stratification and a more marked seasonality with leaf fall, so temperate forest canopies probably shelter a lower proportion of specific taxa (Ulyshen 2011). Canopy functional biodiversity in temperate forests has therefore received relatively little attention to date (Ulyshen 2011). However, the studies conducted in temperate forests (e.g. Bouget et al. 2011; Vodka and Cizek 2013; Plewa et al. 2017) have shown a clear vertical stratification of insect assemblages, just as in tropical forests, with 20 – 40% of all forest insect species strictly associated with canopies (Bouget et al. 2011). In addition to these specialist species, many Arthropods also rely on the canopy for a part of their life cycle, for maturation feeding and mating on foliage, such as *Agrilus* spp. for instance (Ulyshen 2011; Sallé et al. 2014). However, canopies are still relatively unknown biotic frontiers. These crown ecosystems harbor poorly understood, rarely described (both in terms of composition and abundance) insect communities (Bouget et al. 2011). They potentially shelter an underestimated pool of rare or patrimonial species (Plewa et al. 2017), but also native and invasive pests.

Canopy modifications in response to decline may change resource availability and microclimates and may create novel colonization opportunities, thus modulating in different ways the community dynamics of canopy-dwelling insects, depending on their functional guilds. Changes in foliage quality during plant stress may influence the performance of leaf feeders, but the magnitude and orientation of the herbivore response likely depends on both stress intensity and the feeding strategy of the herbivore (Larsson 1989; Herms and Mattson 1992). In addition, the decrease in the number of living branches in the canopy of declining trees may also negatively affect leaf-, seed-, and flower-feeding species (Martel and Mauffette 1997). A survey of Lepidopteran communities in declining maple stands indicated that exposed caterpillars became more abundant while the density of semi-concealed or endophagous species decreased (Martel and Mauffette 1997). This suggests that phyllophagous or seminiphagous insects with an intimate relationship with their host-tree, like specialist species with an endophytic larval development, may be negatively affected by the decrease in foliage quantity or quality and/or the change in microclimate, while these modifications might promote generalist species (Martel and Mauffette, 1997). Conversely, saproxylic beetles are likely to show a marked positive response to forest decline, both in terms of abundance and species richness. Saproxylic beetles form a functional guild associated with dead and decaying wood, related microhabitats, and other saproxylic taxa (Stokland et al. 2012). This guild also includes xylophagous species developing on weakened trees and acting as secondary pests, like the buprestids (Coleoptera: Buprestidae), which are contributing agents during declines (Sallé et al., 2014; Tiberi et al. 2016). The weakened trees and the accumulation of dead wood and related microhabitats typical of declining stands should promote the abundance and diversity of this functional guild.

Our investigation focused on oak forests, which have at least two relevant characteristics for our study purpose. First, oak forests have regularly undergone periods of decline throughout Europe during the last centuries (e.g. Delatour, 1983; Oszako, 2000; Thomas et al., 2002; Sonesson and Drobyshev, 2010; Denman et al., 2014). Moreover, the frequency and intensity of declines have recently increased, and extended canopy modifications have already been documented in Mediterranean oak forests (Allen et al. 2010; Carnicer et al. 2011; Millar and Stephenson 2015). Second, oak forests host a species-rich insect fauna (Southwood 1961). We sampled the communities of leaf-dwelling weevils (Coleoptera: Curculionidae) and saproxylic beetles for three consecutive years in oak stands exhibiting different levels of tree decline.

Firstly, we hypothesized that saproxylic beetles, especially xylophagous species, would be favored by decline. Secondly, we expected to find contrasted responses to decline intensity for leaf-feeding weevils, dependent on their relationship with the host plant. More specifically, we hypothesized that weevils with endophytic larvae would be negatively affected by decline while species feeding on foliage only during the adult stage would be favored. Finally, we hypothesized that seminiphagous weevils would be negatively affected by the reduced amount of acorns in declining stands. Consequently, our objectives were (i) to describe the canopy-dwelling communities of buprestid beetles, other saproxylic species and weevils, and (ii) to evaluate how the local intensity of forest decline was modifying the diversity of these communities.

## Material and methods

### Study area

The study was conducted in the two adjacent state forests of Vierzon and Vouzeron, with a surface area of 5,300 ha and 2,200 ha respectively, located in the center of France 200 km south of Paris (47° 26’ 89” N, 02° 10’ 74” E). The Vierzon forest is dominated by oaks (mostly *Quercus petraea* and *Quercus robur),* at 70%, and conifers (mostly *Pinus sylvestris* and *Pinus nigra),* at 30%, in both pure and mixed stands. The Vouzeron forest is dominated by conifers (mostly *Pinus sylvestris* and *Pinus nigra),* at 65%, in pure stands or mixed with *Q. robur* and *Q. petraea.* The oaks, especially *Q. robur,* in the Vierzon forest have suffered from regular declines (documented in 1920, 1940 and 1982 (Douzon 2006)). The last severe oak decline occurred between 2000 and 2010, which was followed by a sanitation cutting of 100,000 m^3^ of oak over 1000 ha (Douzon, 2006). Several factors were implicated in these successive periods of decline. The prominent predisposing factor was edaphic. In most areas the water table is shallow and variable, and therefore inadequate for the development *Q. robur,* which was however extensively planted in this forest (Douzon 2006; Marçais and Desprez-Loustau 2014). The prominent inciting factors were severe droughts and defoliation caused by powdery mildew (Douzon, 2006; Marçais and Desprez-Loustau 2014). Finally, the most frequently observed contributing biotic agents were *Agrilus biguttatus* Fabricius and *Armillaria mellea* (Vahl ex Fr.) P. Kumm. (Douzon 2006).

### Site selection and characterization

Overall, 14 stands dominated by mature oaks were monitored during our study in the two forests (table 1). In 2016, 13 plots were set up in 11 stands (table 1). Plots were homogeneous areas (approx. 2,000 m^2^) within stands in terms of tree composition and dendrometric parameters. Most plots were located in different stands (table 1). They were located at a minimum of 100 m from the others, but in most cases several kilometers from each other. In 2017 and 2018, 12 plots located in 11 stands were selected in the two forests (table 1). Some of the original plots were changed in 2018 because the stands had been either cleared or harvested selectively (table 1). We selected one tree at the center of each plot, on which we suspended one trap for beetle collection (see below “Beetle sampling”).

**Table 1:**
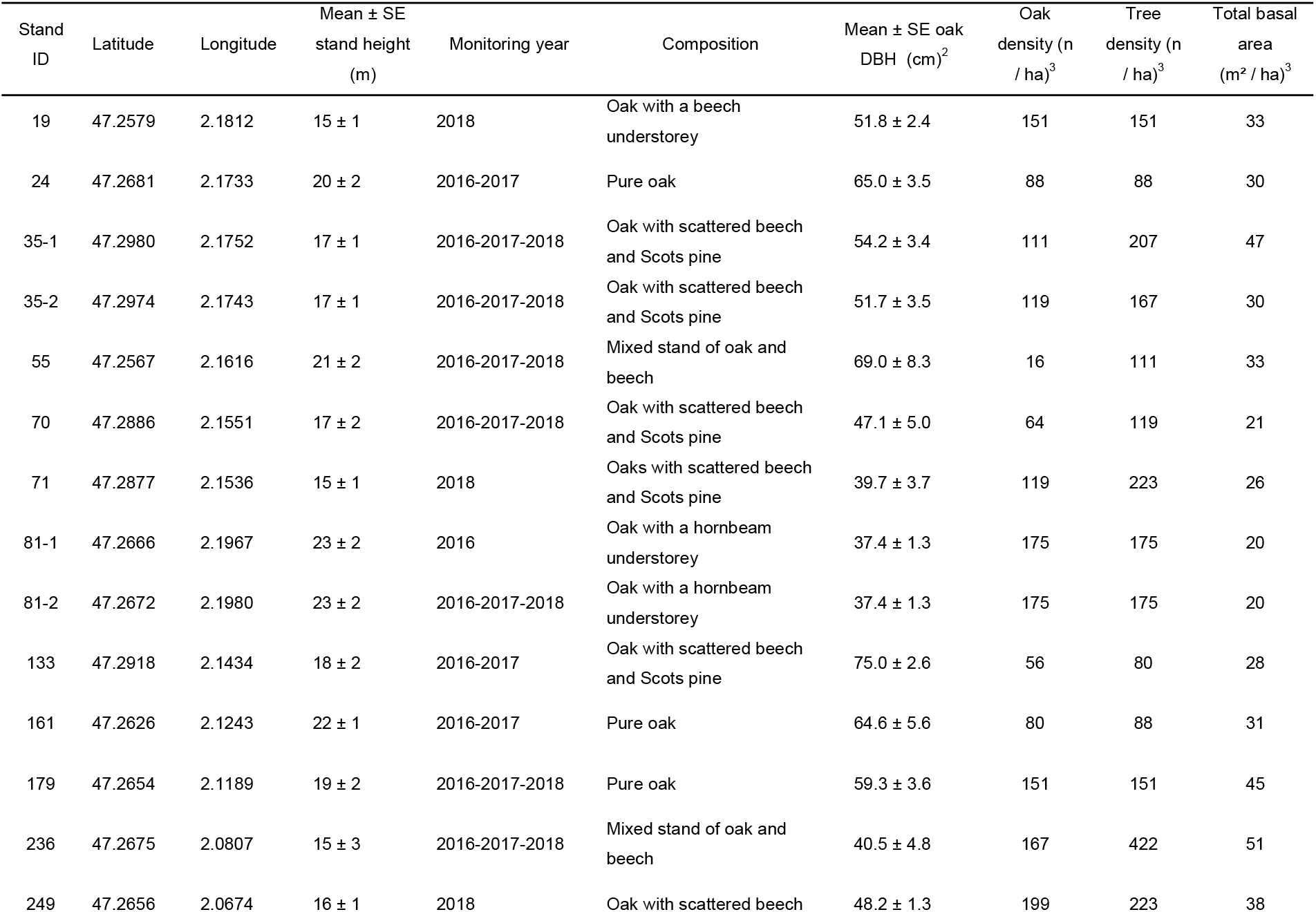
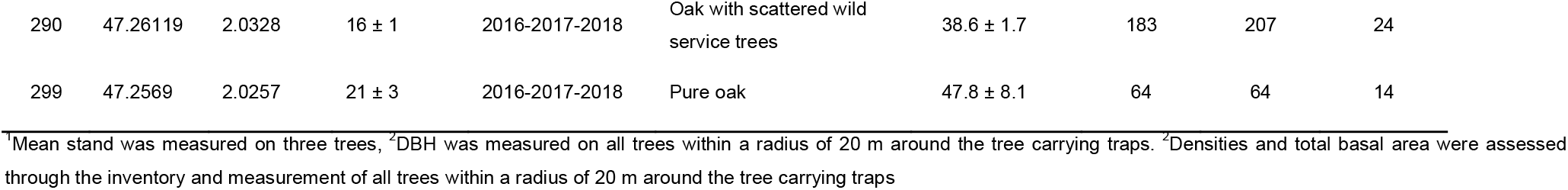
characteristics of the monitored oak stands

| Stand ID | Latitude | Longitude | Mean ± SE stand height (m) | Monitoring year | Composition | Mean ± SE oak DBH (cm)<sup>2</sup> | Oak density (n / ha)<sup>3</sup> | Tree density (n / ha)<sup>3</sup> | Total basal area (m<sup>2</sup> / ha)<sup>3</sup> |
| --- | --- | --- | --- | --- | --- | --- | --- | --- | --- |
| 19 | 47.2579 | 2.1812 | 15 ± 1 | 2018 | Oak with a beech understorey | 51.8 ± 2.4 | 151 | 151 | 33 |
| 24 | 47.2681 | 2.1733 | 20 ± 2 | 2016-2017 | Pure oak | 65.0 ± 3.5 | 88 | 88 | 30 |
| 35-1 | 47.2980 | 2.1752 | 17 ± 1 | 2016-2017-2018 | Oak with scattered beech and Scots pine | 54.2 ± 3.4 | 111 | 207 | 47 |
| 35-2 | 47.2974 | 2.1743 | 17 ± 1 | 2016-2017-2018 | Oak with scattered beech and Scots pine | 51.7 ± 3.5 | 119 | 167 | 30 |
| 55 | 47.2567 | 2.1616 | 21 ± 2 | 2016-2017-2018 | Mixed stand of oak and beech | 69.0 ± 8.3 | 16 | 111 | 33 |
| 70 | 47.2886 | 2.1551 | 17 ± 2 | 2016-2017-2018 | Oak with scattered beech and Scots pine | 47.1 ± 5.0 | 64 | 119 | 21 |
| 71 | 47.2877 | 2.1536 | 15 ± 1 | 2018 | Oaks with scattered beech and Scots pine | 39.7 ± 3.7 | 119 | 223 | 26 |
| 81-1 | 47.2666 | 2.1967 | 23 ± 2 | 2016 | Oak with a hornbeam understorey | 37.4 ± 1.3 | 175 | 175 | 20 |
| 81-2 | 47.2672 | 2.1980 | 23 ± 2 | 2016-2017-2018 | Oak with a hornbeam understorey | 37.4 ± 1.3 | 175 | 175 | 20 |
| 133 | 47.2918 | 2.1434 | 18 ± 2 | 2016-2017 | Oak with scattered beech and Scots pine | 75.0 ± 2.6 | 56 | 80 | 28 |
| 161 | 47.2626 | 2.1243 | 22 ± 1 | 2016-2017 | Pure oak | 64.6 ± 5.6 | 80 | 88 | 31 |
| 179 | 47.2654 | 2.1189 | 19 ± 2 | 2016-2017-2018 | Pure oak | 59.3 ± 3.6 | 151 | 151 | 45 |
| 236 | 47.2675 | 2.0807 | 15 ± 3 | 2016-2017-2018 | Mixed stand of oak and beech | 40.5 ± 4.8 | 167 | 422 | 51 |
| 249 | 47.2656 | 2.0674 | 16 ± 1 | 2018 | Oak with scattered beech | 48.2 ± 1.3 | 199 | 223 | 38 |
| 290 | 47.26119 | 2.0328 | 16 ± 1 | 2016-2017-2018 | Oak with scattered wild<br>service trees | 38.6 ± 1.7 | 183 | 207 | 24 |
| 299 | 47.2569 | 2.0257 | 21 ± 3 | 2016-2017-2018 | Pure oak | 47.8 ± 8.1 | 64 | 64 | 14 |

<sup>1</sup>Mean stand was measured on three trees, <sup>2</sup>DBH was measured on all trees within a radius of 20 m around the tree carrying traps. <sup>3</sup>Densities and total basal area were assessed through the inventory and measurement of all trees within a radius of 20 m around the tree carrying traps

The level of decline was evaluated yearly, during winter, at two embedded spatial scales. (i) At the tree scale: individual decline status was assessed for all bearing-trap trees. (ii) At the plot scale: decline status was assessed for trees surrounding the bearing-trap tree, which encompassed only the five closest oak trees in 2016 and 2017, and all the oaks located within a radius of 20 m around the bearing-trap tree in 2018. Decline level was evaluated following the protocol designed by the French Forest Health Service (Département de la Santé des Forêts, DSF) (Nageleisen 2005). In brief, crown transparency, the proportion of branches without leaves, the proportion of dead branches and leaf distribution in the canopy were evaluated. Based on these criteria, each tree was given a decline index ranging from 0 (no decline) to 5 (dead tree) (table 2). Trees with an index value equal to or below two were considered healthy. Trees with an index value equal to or above three were considered in decline. For the plot scale, the proportion of declining trees (with a decline index equal to or above three) was calculated, following a routine DSF procedure (table 2).

**Table 2:** decline index of trees carrying traps and percentage of surrounding declining trees in the monitored oak plots, for the three survey years

| Stand<br>ID | Decline index of trees with traps |  |  | Percentage of surrounding declining trees |  |  |
| --- | --- | --- | --- | --- | --- | --- |
|  | 2016 | 2017 | 2018 | 2016 | 2017 | 2018 |
| 19 | NA | NA | 3 | NA | NA | 89 |
| 24 | 4 | 3 | NA | 17 | 33 | NA |
| 35-1 | 2 | 1 | 1 | 17 | 0 | 19 |
| 35-2 | 0 | 1 | 1 | 17 | 33 | 14 |
| 55 | 3 | 3 | 3 | 83 | 67 | 100 |
| 70 | 3 | 3 | 3 | 83 | 71 | 100 |
| 71 | NA | NA | 3 | NA | NA | 87 |
| 81-1 | 1 | 1 | 1 | 17 | 17 | 14 |
| 81-2 | 2 | NA | NA | 0 | NA | NA |
| 133 | 3.5 | 2 | NA | 33 | 57 | NA |
| 161 | 2 | 2 | NA | 33 | 29 | NA |
| 179 | 2 | 4 | 2 | 17 | 57 | 61 |
| 236 | 3 | 3 | 3 | 33 | 33 | 45 |
| 249 | NA | NA | 3.5 | NA | NA | 60 |
| 290 | 2 | 2 | 2 | 33 | 14 | 43 |
| 299 | 3.5 | 2 | 2 | 50 | 57 | 50 |

### Beetle sampling

#### Optimization of the sampling protocol

We used multi-funnel traps (Lindgren traps, Chemtica Internacional, San Jose, Costa Rica), each with 12 fluon-coated funnels, to collect the insects. To optimize our protocol for sampling canopy-dwelling beetles, we tested two trap heights and two trap colors. In 2016, we assessed whether trap height significantly affected the composition or relative abundance of the captured species. To do so, we suspended two green traps at different heights in the same tree, one approximately 15 m from the ground and the other 10 m from the ground. Thirteen pairs of traps were considered.

In 2017 and 2018, we also compared the trapping efficiency of green and purple multi-funnel traps. Green multi-funnel traps have successfully captured a wide array of buprestid species in North America and Europe (Petrice and Haack 2015; Rassati et al. 2019), but Brown et al. (2017) showed that some European *Agrilus* species might be more attracted to purple than to green. In ten trees, one green and one purple trap were both suspended roughly 15 m above the ground.

#### Routine sampling protocol

For the sake of consistency, and because it was the best sampling design (see “Results), only green multi-funnel traps, suspended at 15 m from the ground (i.e., among the lower branches of the canopy) were considered to assess the effect of decline on canopy-dwelling beetles. One trap was set up within each monitored plot. The collectors were filled with a 50% (v/v) monopropylene glycol solution diluted with water, and a drop of detergent. No lure was added to the traps. The traps were emptied every month and the captured species were recorded. In 2016, the traps were installed in June and collection continued until September. In 2017, the traps were installed in April and collection continued until October. In 2018, the traps were installed in April and collection continued until September.

#### Beetle identification and ecological trait assignment

Three beetle groups were considered for analysis: (i) oak-associated buprestid beetles, i.e. xylophagous species specifically attracted by green Lindgren traps; (ii) other saproxylic beetles, excluding “tourist” species associated to conifer tree species, split into two feeding guilds, i.e. the xylophagous species guild (incl. xylophagous *sensu stricto* and saproxylophagous species) and the non-xylophagous species guild (incl. saprophagous, zoophagous, mycetophagous species); and (iii) oak-associated phytophagous weevils, which were split into two feeding guilds, i.e. the leaf-eating (phyllophagous, i.e. folivore) species guild and the seed- and fruit-eating (seminiphagous, i.e. acorn borers) species guild. In the phyllophagous guild, we considered species whose adults feed on oak leaves (mainly leaf chewers) as generalist species, and species whose both larvae and adults feed on oak leaves (larvae dwelling in foliar tissues) as specialist species.

Certain saproxylic families were removed from the data set (Latridiidae, Leiodidae, Malachiidae, Cantharidae, Corylophidae, Cryptophagidae, Ptiliidae, Staphylinidae), because they are often difficult to identify at the species level and due to the lack of specialists able to check species identifications. The French Frisbee database was used as the reference list of feeding guilds for saproxylic beetle species from the 39 recorded beetle families (Bouget et al. 2019). Most of the beetle specimens were identified by several of the authors (GP, BN, XP, RB, TFG and RL). The remaining families were identified by specialists, as mentioned in the Acknowledgements. For each guild, we computed number of species (richness), number of individuals (abundance) and beetle biomass caught at the trap level and cumulated over all the sampling season per year. Biomass, actually dry weight (in mg), was assessed through the following formula: Biomass = 3.269+L^2.463^, where “L” is the body length in millimeters (Ganihar 1997 in Seibold et al. 2019). The abundance of several dominant species of oak-associated buprestids and phytophagous weevils was also analyzed.

#### Data analysis

All analyses were performed in R, version 3.5.1 (R Core Team 2018). Trap color and trap height effects on catches of buprestids, other saproxylic beetles and phytophagous weevils were assessed with Wilcoxon signed-ranked tests. To analyze the effect of the decline level at tree scale on cumulative number of detected species, we rarefied species richness to the same sample size (interpolated rarefaction sampling without replacement; specaccum function, vegan R-package). To rank the effect of decline level at tree or plot scale on variations in average univariate metrics (mean values per trap of guild richness, species abundance, guild abundance, guild biomass), we used the differences in the Akaike information criterion (AICc) scores to compare the fit between the generalized linear mixed models including separately each of the two explanatory variables and their fit with the null model. To assess the significance of the estimates of the best decline features for each response variable, the error structure of the generalized linear mixed-effects models was adjusted to better fit the data. To do so, glmm were fitted for the negative binomial family, the Gaussian family, the log-normal family (i.e. log-transformed response), and the Poisson family (functions glmer.nb, glmer, lmer, lme4 R-package). To account for repeated measures and configuration of sampling design, plot and year were added as nested random effects on the intercept in mixed models. To rank the effect of decline level at tree or tree-group scale on variations in species composition (including singletons), we performed a Canonical Analysis of Principal coordinates (function capscale, vegan R-package, CAP, Anderson and Willis 2003). Based on Bray-Curtis distance matrices, we carried out inertia partitioning on all the explanatory environmental variables, as colinearity among predictor variables is not a problem in CAP. We calculated total constrained inertia, the total inertia explained by each variable, the latter’s statistical significance (permutation tests – 100 runs) and the relative individual contribution of each variable to constrained inertia.

We used the IndVal method to identify beetle species indicating tree decline level (healthy *vs.* declining) (Dufrêne and Legendre 1997, indicspecies R-package). This method calculates the association value (IndVal index) between the species and a group of sites, based on between-group variations in occurrence (fidelity) and abundance (specificity), and then tests the statistical significance of this relationship with a permutation test. P-values were corrected for multiple testing. Only species shown to be significant in the permutation test with an indicator value above 25%, occurring in more than 10% of the samples and with more than 10 individuals sampled were considered here.

## Results

For all guilds, the number of individuals captured was higher in the upper traps (Fig. 1). For both guilds of leaf-dwelling species (i.e. Agrilinae and phytophagous weevils), green traps were markedly more attractive than purple traps, while no difference between traps was detected for the other saproxylic species (Fig. 2).

**Figure 1:**
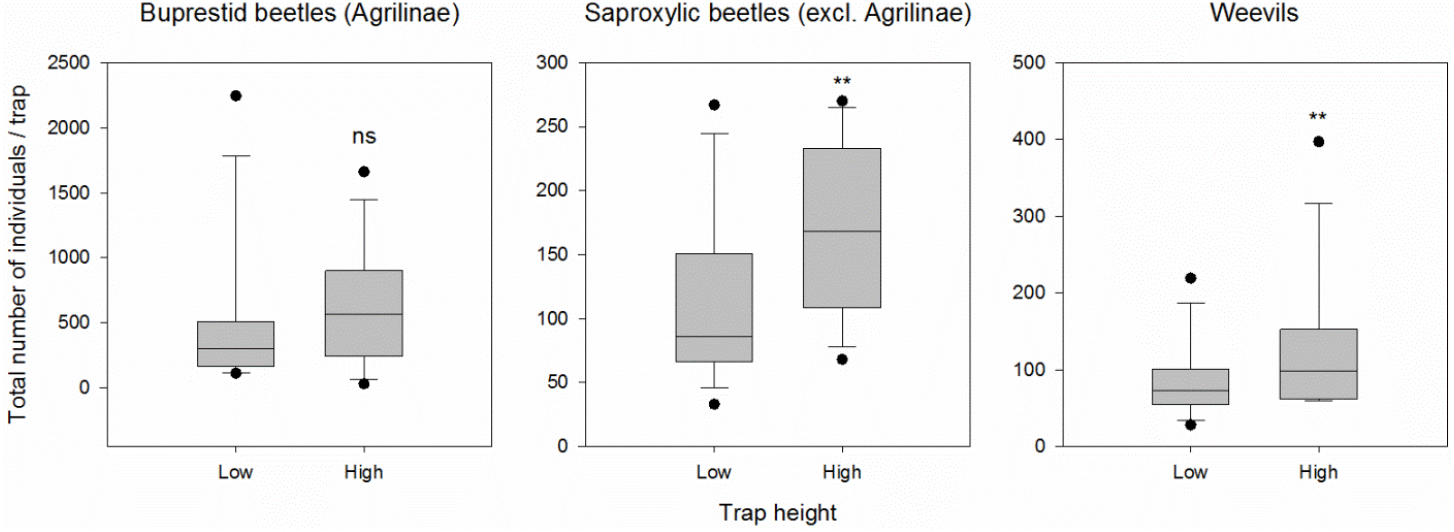
effect of trap height (10 m *vs.* 15 m above the ground) on the number of oak-associated Agrilinae (i.e. *Agrilus* sp., *Coraebus* sp. and *Meliboeus* sp.), other saproxylic beetles, and phytophagous weevils (i.e. phyllophagous and seminiphagous species) captured per trap. P<0.01:**.

**Figure 2.**
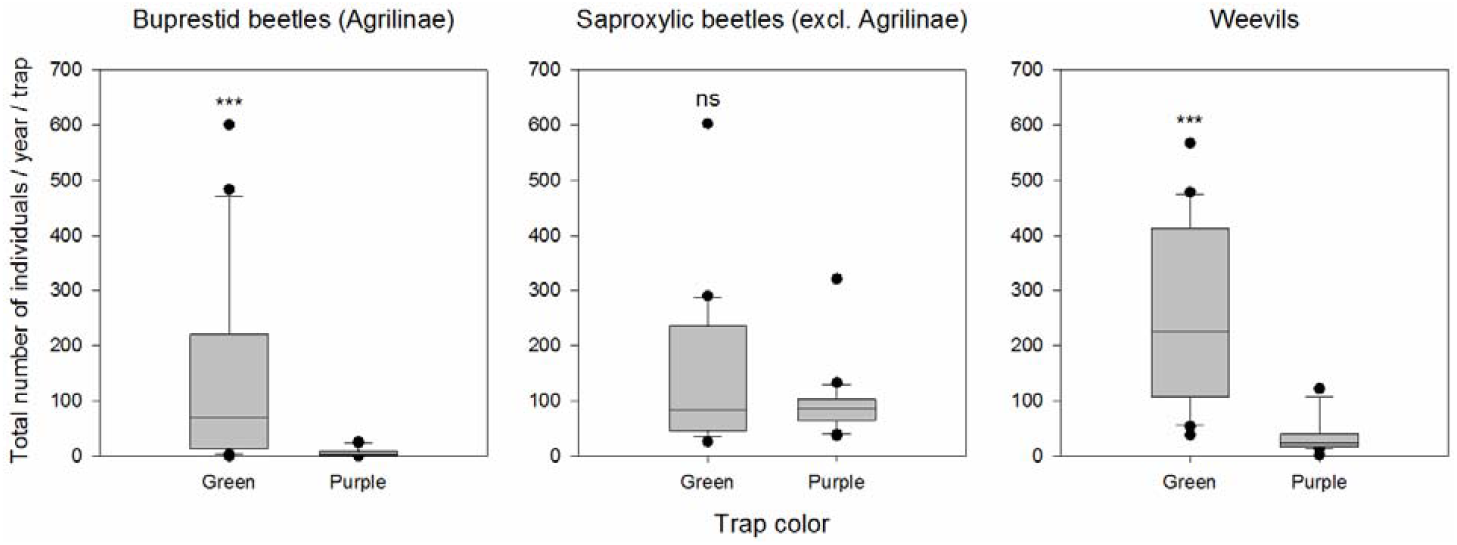
Effect of trap color (green *vs.* purple) on the number of oak-associated Agrilinae (i.e. *Agrilus* sp., *Coraebus* sp. and *Meliboeus* sp.), other saproxylic beetles, and phytophagous weevils (i.e. phyllophagous and seminiphagous species) captured per trap. P<0.01: **, P<0.001: ***.

Overall, for the assessment of decline effects on canopy-dwelling beetles, the compiled data set of 27,627 individual specimens included 266 beetle species: 10,440 individuals and ten species of oak-associated buprestid beetles; 8,280 individuals and 21 species of oak-associated phytophagous weevils (4 seminiphagous, 10 specialist phyllophagous, 5 generalist phyllophagous and 2 flower-eating (anthophagous) species); 3,008 individuals and 102 species of non-xylophagous saproxylic beetles; and 5,899 individuals and 133 species of xylophagous saproxylic beetles (table S1). This corresponds to 14,490 individuals and 223 beetle species found in the 23 traps hanging from non- or slightly-declining trees, and 13,137 individuals and 194 species found in the 14 traps hanging from declining trees. On the whole, cumulative species-richness estimates at a standardized sample size did not display any significant contrast between decline levels at the tree scale, either for the whole beetle community, or for individual guilds (Fig. S1).

We detected many significant effects of decline level on guild metrics (mean abundance, biomass and richness per trap) and on species mean abundances (table 3). Most of these effects were positive, except, at the plot scale, for (i) a negative effect of decline level on the mean abundance of xylophagous beetles (table 3 and Fig. 3), and (ii) a negative effect of decline level on the mean abundance of two specialist phyllophagous species, i.e. *Archarius pyrrhoceras* Marsham and *Orchestes quercus* L. (table 3). We measured significant positive effects of decline level at the tree scale on the species richness of xylophagous beetles (Fig. 3) and on the biomass and abundance of buprestids (Fig. 4); and at plot scale, on the biomass of non-xylophagous saproxylic beetles (Fig. 3), on species richness of buprestids (Fig. 4) and phyllophagous weevils (Fig. 5), and on the abundance of generalist phyllophagous weevils (Fig. 5). Five of the six buprestid species tested responded positively in abundance to decline intensity (at tree scale: *Agrilus angustulus* Illiger, *A. biguttatus, Agrilus laticornis* Illiger, *Agrilus obscuricollis* Kiesenwetter, *Agrilus sulcicollis* Lacordaire; and at the plot scale: *Coraebus undatus* Fabricius), as well as one of the two tested generalist phyllophagous weevil *(Phyllobius pyri* L.) (table 3). Seminiphagous species were not significantly affected by decline level at any scale, either at species or guild level (table 3, Fig. 5). Specialist phyllophagous weevils responded to decline intensity at the species level but not at the guild level (table 3, Fig. 5). When all the sampled species were pooled, we also observed significant positive effects of decline at the tree scale on the biomass and abundance of all beetles (table 3, Fig. 6).

**Figure 3:**
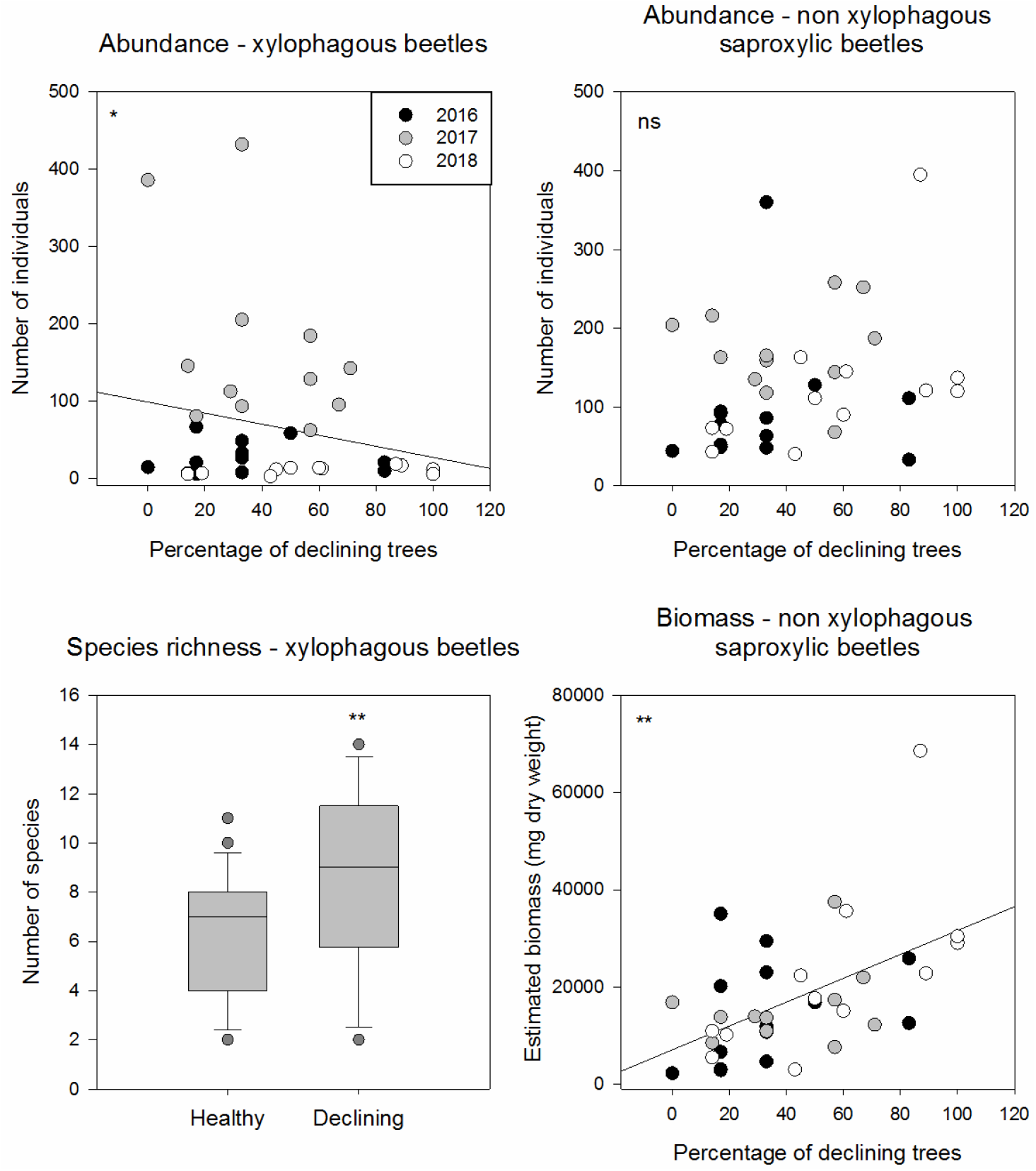
abundance of xylophagous and non-xylophagous saproxylic beetles, species richness of xylophagous beetles, and biomass of non-xylophagous saproxylic beetles depending on decline level at the tree scale (heatlhy vs. declining) or plot scale (percentage of declining trees). See table 3 for statistical results; P<0.05: *; P<0.01: **.

**Figure 4:**
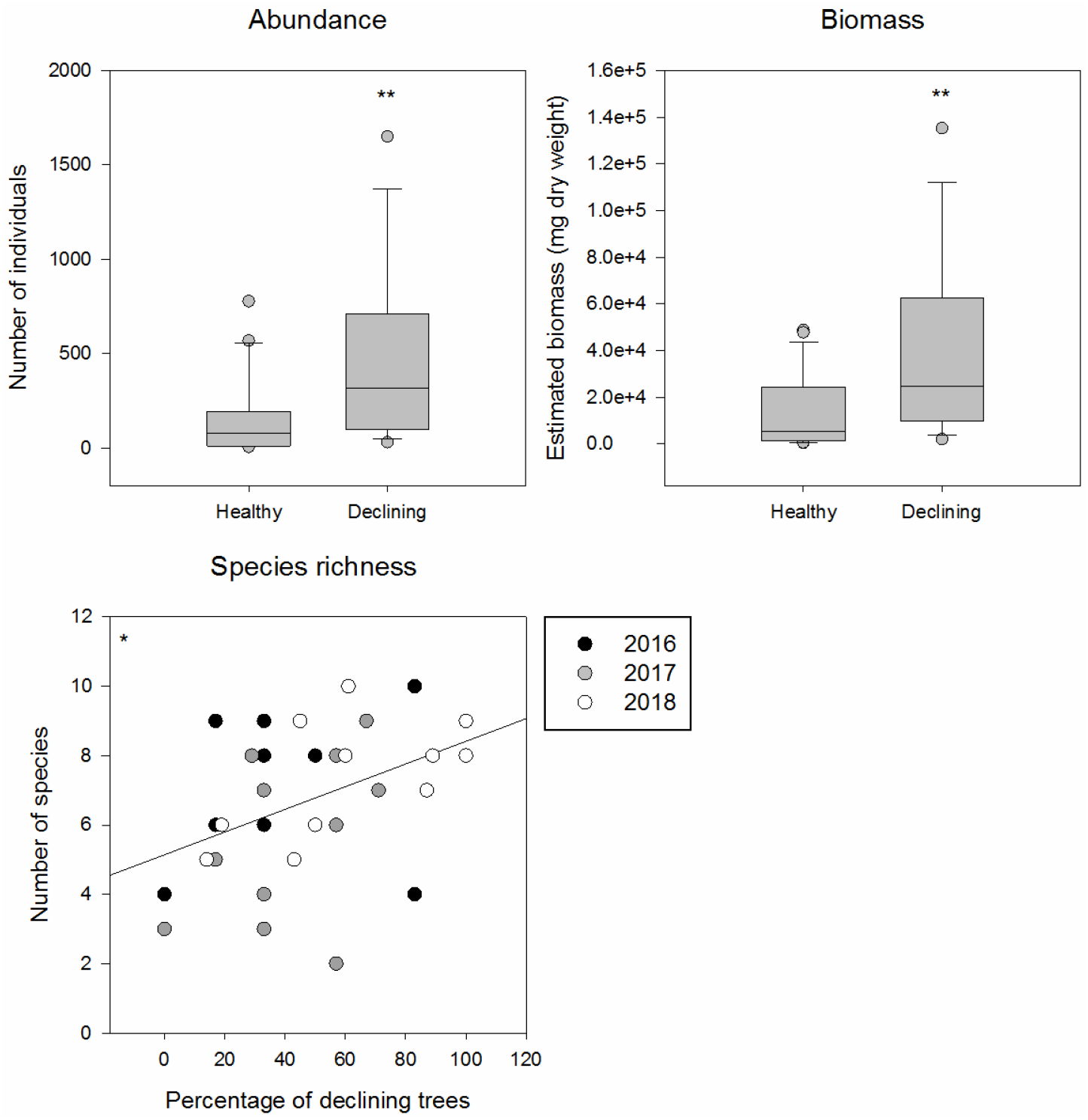
abundance, biomass and species richness of oak-associated buprestid beetles depending on decline level at the tree scale (heatlhy vs. declining) or plot scale (percentage of declining trees). See table 3 for statistical results; P<0.05: *; P<0.01: **.

**Figure 5:**
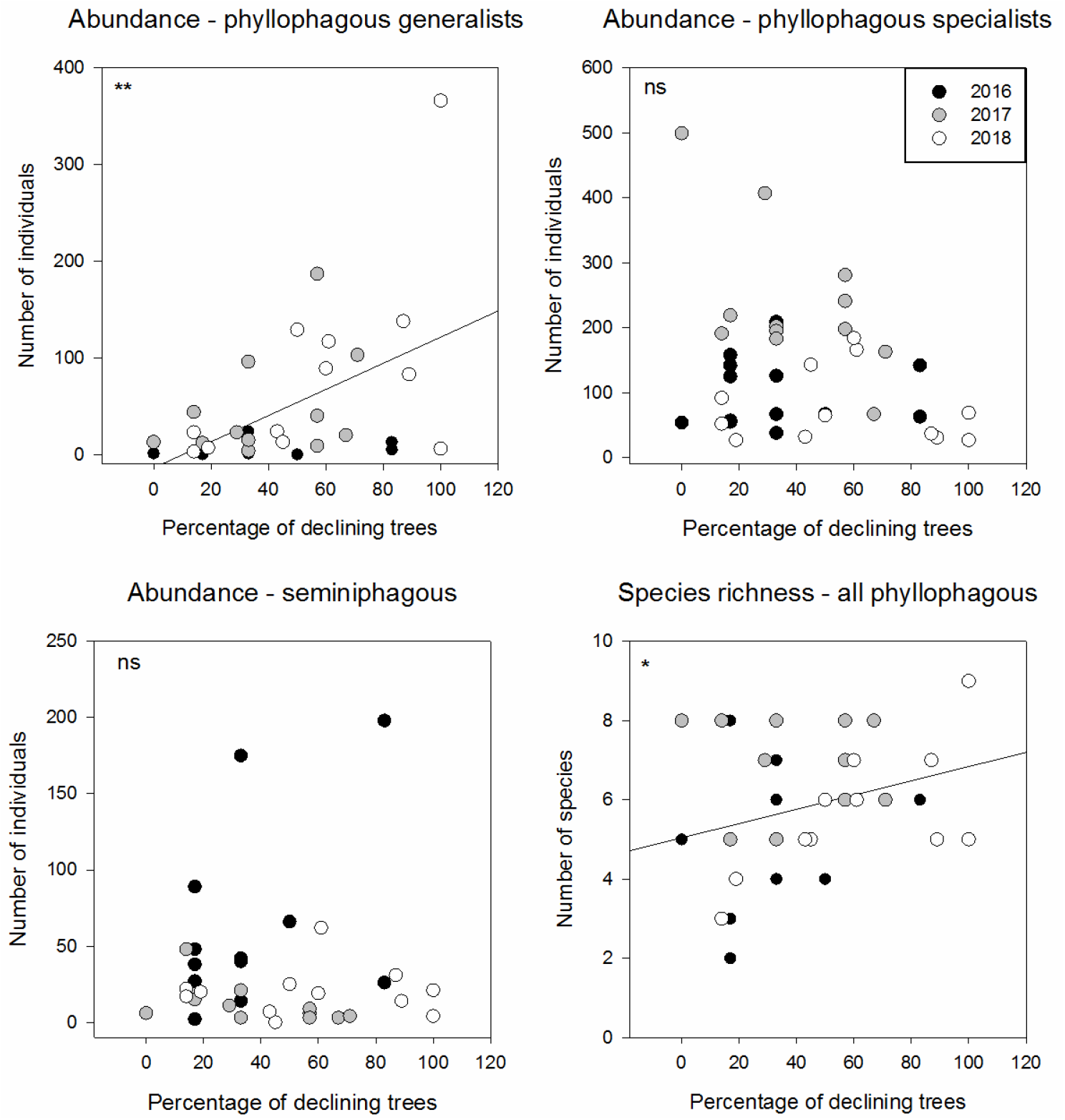
abundance of generalist phyllophagous, specialist phyllophagous and seminiphagous weevils, and species richness of all phyllophagous weevils, depending on decline level at the plot scale (percentage of declining trees). See table 3 for statistical results; P<0.05: *; P<0.01: **.

**Figure 6:**
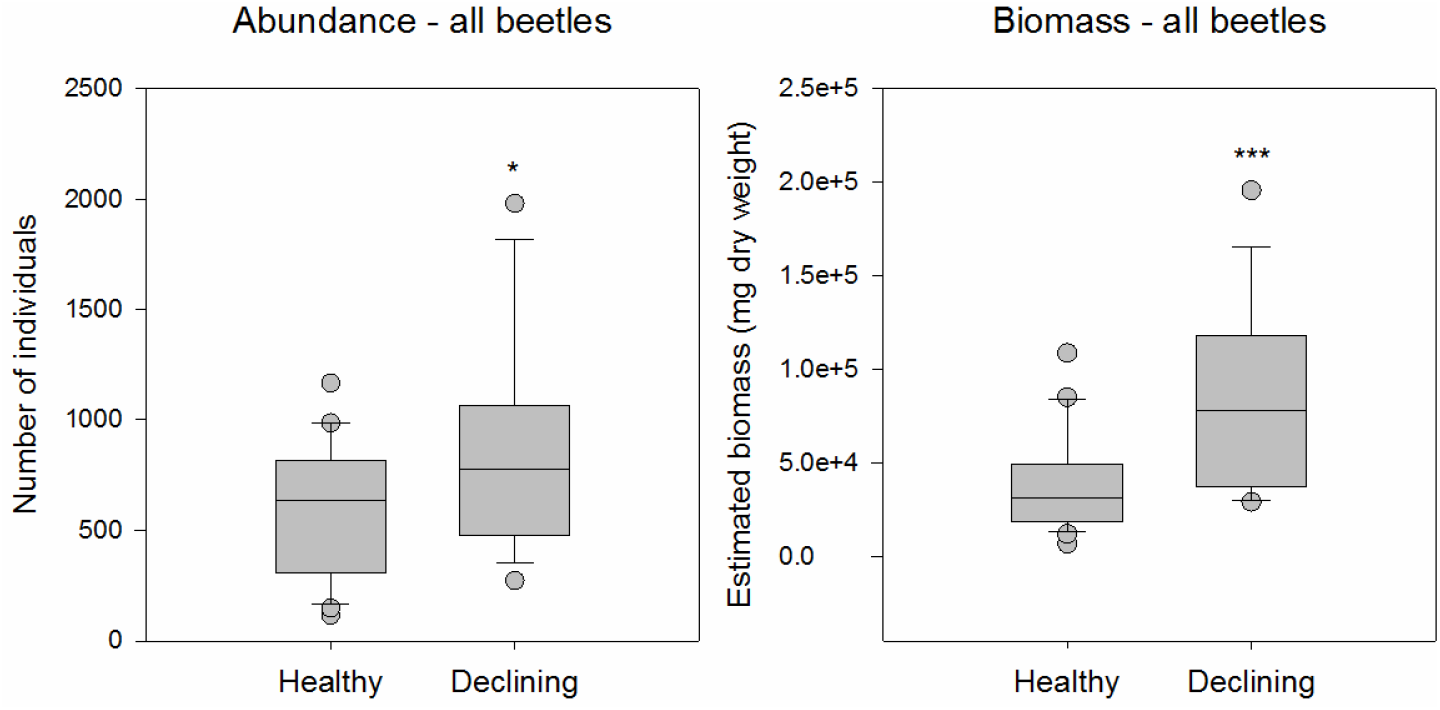
abundance and biomass of all the beetle species considered in the analyses, depending on decline at the tree scale (heatlhy vs. declining). See table 3 for statistical results; P<0.05: *; P<0.01: **.

**Table 3:** effect of decline on biomass, abundance and species richness for the different groups and guilds of beetles, and on abundance for the main species collected in the canopy.

|  |  | Variable<br>(mean value per trap) | Best ecological model | Delta[AICc]<sup>°</sup> | Effect of decline<br>estimate | SE |
| --- | --- | --- | --- | --- | --- | --- |
| Oak-associated buprestid beetles |  | biomass<sup>(3)</sup> | Tree | -6 | **1.26** ** | 0.44 |
|  |  | abundance<sup>(1)</sup> | Tree | -5 | **1.38** ** | 0.48 |
|  |  | species richness<sup>(4)</sup> | Plot | -2 | **0.14** * | 0.06 |
|  |  | *Agrilus angustulus*<sup>(3)°°</sup> | Tree | -4 | **1.49** * | 0.59 |
|  |  | *Agrilus biguttatus*<sup>(1)</sup> | Tree | -2 | **2.03** * | 0.93 |
|  |  | *Agrilus hastulifer*<sup>(1)</sup> | Tree | 0 | 1.84 | 1.05 |
|  |  | *Agrilus laticornis*<sup>(1)</sup> | Tree | -2 | **0.98** * | 0.46 |
|  |  | *Agrilus obscuricollis*<sup>(1)</sup> | Tree | -2 | **1.87** * | 0.93 |
|  |  | *Agrilus sulcicollis*<sup>(1)</sup> | Tree | -9 | **1.88** *** | 0.51 |
|  |  | *Coraeus undatus*<sup>(1)</sup> | Plot | -7 | **0.94** *** | 0.28 |
| Saproxylic beetles (excl. buprestids) | Xylophagous | biomass<sup>(3)</sup> | Tree | 0 | 0.71 | 0.46 |
|  |  | abundance<sup>(3)</sup> | Plot | -2 | **-0.39** * | 0.16 |
|  |  | species richness<sup>(1)</sup> | Tree | -1 | **0.29** * | 0.14 |
|  | Non-xylophagous | biomass<sup>(3)</sup> | Plot | -3 | **0.31** ** | 0.10 |
|  |  | abundance<sup>(1)</sup> | Plot | +1 | 0.12 | 0.10 |
|  |  | species richness<sup>(2)</sup> | Plot | 0 | 3.41 | 1.84 |
| Oak-associated weevils | All phytophagous | biomass<sup>(3)</sup> | Plot | 0 | 0.19 | 0.12 |
|  |  | abundance<sup>(3)</sup> | Tree | +2 | 0.10 | 0.10 |
|  |  | species richness<sup>(2)</sup> | Plot | 0 | 0.64 | 0.37 |
|  | All phyllophagous | biomass<sup>(3)</sup> | Tree | -1 | 0.23 | 0.13 |
|  |  | abundance<sup>(3)</sup> | Tree | +2 | -0.10 | 0.26 |
|  |  | species richness<sup>(3)</sup> | Plot | -3 | **0.10** * | 0.04 |
|  | Generalist phyllophagous | biomass<sup>(3)</sup> | Plot | -1 | 0.59 | 0.32 |
|  |  | abundance<sup>(3)</sup> | Plot | -5 | **0.77** ** | 0.23 |
|  |  | species richness<sup>(3)</sup> | Tree | +5 | 0.04 | 0.12 |
|  |  | *Phyllobius pyri*<sup>(1)</sup> | Plot | -6 | **1.69** ** | 0.56 |
|  |  | *Polydrusus cervinus*<sup>(1)</sup> | Plot | +2 | 0.27 | 0.28 |
|  | Specialist phyllophagous | biomass<sup>(3)</sup> | Tree | +2 | -0.10 | 0.04 |
|  |  | abundance<sup>(3)</sup> | Tree | +1 | -0.12 | 0.10 |
|  |  | species richness<sup>(4)</sup> | Plot | 0 | 0.14 | 0.09 |
|  |  | *Archarius pyrrhoceras*<sup>(1)</sup> | Plot | -3 | **-0.79** * | 0.37 |
|  |  | *Orchestes quercus*<sup>(1)</sup> | Plot | -1 | **-0.27** * | 0.13 |
|  | Seminiphagous | biomass<sup>(3)</sup> | Tree | +1 | -0.38 | 0.57 |
|  |  | abundance<sup>(1)</sup> | Plot | +2 | -0.14 | 0.21 |
|  |  | species richness<sup>(4)</sup> | Plot | +2 | 0.06 | 0.10 |
|  |  | *Curculio glandium*<sup>(1)</sup> | Plot | +2 | -0.15 | 0.22 |
| All beetles |  | biomass<sup>(3)</sup> | Tree | -10 | **0.80** *** | 0.22 |
|  |  | abundance<sup>(2)</sup> | Tree | -3 | **298.2** * | 121.10 |
|  |  | species richness<sup>(2)</sup> | Tree | +1 | 3.70 | 2.33 |

Generalized linear mixed-effects models fitted for the negative binomial family<sup>(1)</sup>, the Gaussian family<sup>(2)</sup>, the log-normal family<sup>(3)</sup> (i.e. log-transformed response), the Poisson family<sup>(4)</sup> with year as a random effect ; biomass = dry weight (in mg); ° ΔAICc = AICc (best ecological model) – AICc (null model); °° the variable considered for species is abundance. P<0.05: \*; P<0.01: \*\*; P<0.001: \*\*\*.

From CAP analyses, we estimated low but significant contributions of decline level to variations in the community composition of most of the beetle groups, i.e. phytophagous weevils, buprestids and xylophagous saproxylic, though not for the non-xylophagous saproxylic guild (table 4). Significant effects on community composition were mainly related to decline level at the plot scale, except for xylophagous saproxylic beetles, which were affected at the tree scale. A larger portion of inertia was explained by sampling year.

**Table 4:** Canonical Analysis of Principal coordinates, based on Bray-Curtis distance matrices, ranking the effect of the two spatial levels of decline (plot vs. tree) on variations in species composition.

| Group | Ecological variable<br>with the best<br>contribution to inertia | Inertia explained by the<br>best ecological variable<br>(and significance) | % inertia<br>explained | Inertia explained<br>by sampling year | % inertia explained<br>by sampling year |
| --- | --- | --- | --- | --- | --- |
| Oak-associated<br>buprestid beetles | Plot | **0.60 \*** | 5.9 | **2.19 \*\*\*** | 21.5 |
| Xylophagous beetles<br>(excl. buprestids) | Tree | **0.58 \*** | 6.7 | **4.70 \*\*\*** | 54.5 |
| Non-xylophagous<br>saproxylic beetles | Plot | 0.13 | 1.3 | **3.14 \*\*\*** | 30.4 |
| Oak-associated<br>weevils | Plot | **0.32 \*** | 4.9 | **1.75 \*\*\*** | 26.2 |
| All beetles | Plot | **0.48 \*\*** | 5.5 | **2.63 \*\*\*** | 30.5 |

P<0.001: \*\*\*.

The IndVal analysis detected 15 characteristic species in declining trees, and only one in healthy trees (table 5). The group of species associated with declining trees consisted primarily of xylophagous species, including four species of *Agrilus,* five other xylophagous beetle species, and two saproxylophagous species.

**Table 5:** characteristic species for each tree-decline level, identified using the IndVal approach. We retained only those species significant in the permutation test with an indicator value above 0.25, sampled in more than 10% of traps and with more than 10 individuals.

| Group | Indicator species | Feeding guild | Indicator value | Frequency (%) |
| --- | --- | --- | --- | --- |
| Declining trees | *Agrilus angustulus* | Xylophagous | 0.891 * | 86 |
|  | *Agrilus sulcicollis* | Xylophagous | 0.889 ** | 81 |
|  | *Agrilus hastulifer* | Xylophagous | 0.798 * | 65 |
|  | *Agrilus biguttatus* | Xylophagous | 0.794 ** | 51 |
|  | *Trichoferus pallidus* | Xylophagous | 0.843 ** | 65 |
|  | *Rhagium sycophanta* | Xylophagous | 0.573 * | 16 |
|  | *Xylotrechus antilope* | Xylophagous | 0.622 * | 30 |
|  | *Scolytus intricatus* | Xylophagous | 0.683 ** | 30 |
|  | *Gasterocercus depressirostris* | Xylophagous | 0.517 * | 14 |
|  | *Mordella brachyura* | Saproxylophagous | 0.804 * | 57 |
|  | *Cetonia aurata* | Saproxylophagous | 0.699 ** | 35 |
|  | *Opilo mollis* | Zoophagous | 0.787 * | 62 |
|  | *Stenagostus rhombeus* | Zoophagous | 0.733 * | 59 |
|  | *Lygistopterus sanguineus* | Zoophagous | 0.716 ** | 43 |
| Healthy trees | *Calambus bipustulatus* | Zoophagous | 0.693 ** | 35 |

P<0.05: \*; P<0.01: \*\*.

## Discussion

Oak decline affected the communities of the canopy-dwelling beetles considered in our study differently depending on their feeding guild and/or host specialization. As predicted, the abundance, biomass, and species richness of oak-associated buprestids increased with the decline severity. The abundance of most major Agrilinae species followed a similar pattern. Consequently, most species contributed to this overall increase. Agrilinae preferentially colonize weakened hosts (e.g. Moraal and Hilszczanski 2000; Jennings et al. 2014; Petrice and Haack 2014; Poole et al. 2019). Their abundance is positively influenced by the availability of fresh snags and coarse woody debris in the environment (Redilla and McCullough 2017), and damaged trees (Brück-Dyckhoff et al. 2019); these features typically occur in declining stands. Several of the species collected, namely *A. biguttatus, A. sulcicollis, A. angustulus,* and *C. undatus,* can act as major contributing agents during oak declines (Sallé et al. 2014). Consequently, they may also have exerted a positive feedback by further weakening their host trees, thus contributing to the accumulation of favorable breeding material. Interestingly, three of the four, *A. biguttatus, A. sulcicollis* and *A. angustulus,* were also good indicators of declining trees, together with *Scolytus intricatus* Ratzeburg and *Gasterocercus depressirostris* Fabricius, other contributing agents of oak decline (Saintonge and Nageleisen 2001; Sallé et al. 2014).

However, while the species richness of other xylophagous species also increased in declining stands, their abundance slightly decreased. This might reflect increased competition among xylophagous species in declining stands. However, the variation in abundance of these other xylophagous beetles should be considered with caution since it was mostly driven by variations in the abundance of *Anisandrus dispar* Fabricius. The abundance of this generalist ambrosia beetle might have been loosely connected to oak decline level. The increased availability of resources and habitats in declining stands, especially large woody debris, probably also participated in the observed increase in the biomass of non-xylophagous saproxylic beetles, since the size of saproxylic species tends to increase with the diameter of the available deadwood resources (Brin et al. 2011).

Feeding guilds of phytophagous weevils responded differently to decline severity. The abundance of generalist phyllophagous species, especially *P. pyri,* increased with decline intensity, while the abundance of the two main specialist phyllophagous species, i.e. *O. quercus* and *A. pyrrhoceras,* decreased. These variations support our hypotheses and are congruent with previous observations by Martel and Mauffette (1997) for Lepidopteran communities colonizing maple foliage. They are nonetheless inconsistent with predictions from the insect performance hypothesis by Larsson (1989) concerning the response of folivores with various feeding habits to tree stress. Several factors may have affected the abundance of phyllophagous weevils differently. Environmental constraints can have contrasted effects on both biochemical and morphological leaf traits (Günthardt-Goerg et al. 2013; Hu et al. 2013). Likewise, the greater exposure of leaves in the opened canopies of declining oaks can alter their phytochemical profile, and may have increased their content in phenolic compounds (Yamasaki and Kikuzawa 2003; Lämke and Unsicker 2018). Such modifications may in turn have different impacts on phytophagous insects depending on their feeding guild and specialization (e.g. Gutbrodt et al. 2011; Forkner et al. 2014). In addition to modifying trophic resources or habitats, crown thinning during a decline can also directly impact larval development by altering the thermal buffering provided by the canopy (Martel and Mauffette 1997; Hardwick et al. 2015; De Frenne et al. 2019). Greater leaf exposure will affect leaf microclimate and may have detrimental effects on endophytic larvae (e.g. Pincebourde et al. 2007), like those of *O. quercus* or *A. pyrrhoceras.* Canopy thinning can also affect forest soil microclimates (De Frenne et al. 2013), allowing free-living larvae like those of *P. pyri* to find optimal microclimatic conditions more easily. Finally, greater leaf exposure may also lead to greater predation pressure on leaf-dwelling endophytic larvae (e.g. Tschanz et al. 2005), which would further explain why specialist phyllophagous weevils with endophytic larvae were negatively affected by oak decline. Overall, the negative response of specialist phyllophagous species to decline may relate to a decrease in leaf suitability (phytochemical profile and microclimatic conditions), an increase in predation pressure or reduced food availability (fewer leaves) (Gely et al. 2019). Conversely, the positive response of generalist phyllophagous species to decline severity could stem from a decrease in interspecific competition with decreasing populations of specialist species (Kaplan and Denno 2007), and from improved conditions for larval development. For seminiphagous weevils, no effect of decline was detected, suggesting either that acorn quantity or quality was not markedly affected by oak decline or that the modifications were not significant enough to impact the species considered.

We considered decline level at two spatial scales: the tree and the plot. Overall, we observed significant responses mainly at the tree scale for xylophagous beetles, including oak-associated buprestids, and mainly at the plot scale for phytophagous and non-xylophagous saproxylic beetles. This might reflect differences among guilds in their dispersal capacity and host-selection behavior. For instance, some xylophagous species might have emerged from the declining trees carrying traps or might have been visually and/or chemically attracted by these declining trees, since weakened hosts often attract secondary pests (e.g., Haack and Benjamin 1982). More specifically host volatiles such as terpenes or ethanol emitted by weakened trees can be used by these insects to discriminate suitable hosts (e.g., Montgomery and Vargo 1983; Sánchez-Osorio et al. 2019).

For all the communities we monitored, further experiments would be necessary to identify the main drivers of the variations observed. More specifically, the effect of decline on the abundance of microhabitats and resources such as dead wood, cavities, opportunistic fungi, and acorns should be quantified (Heitzman et al. 2007; Spetich 2007). Likewise, changes in microclimates and predation pressure at the canopy and soil levels during a decline should be characterized. In addition, in our study we were not able to take into account decline dynamics, since historical data on decline onset, duration and intensity at the stand scale was lacking. We considered stands exhibiting different decline levels, which may result from disturbances with different frequency, severity and/or spatial and temporal extents at the stand scale. Past disturbance regimes can modulate the current taxonomic, functional and phylogenetic composition forest communities, notably the community of saproxylic beetles (Kozák et al. 2020). Therefore integrating historical data in future studies would help to disentangle current decline effects from past disturbance legacies.

Changes in species richness and abundance led to significant community modifications for both xylophagous beetles and phytophagous weevils, which in turn contributed to a significant modification of the overall beetle community. From a functional standpoint, this type of modification may modulate important processes in forest ecosystems, since saproxylic insects play a significant role in wood decomposition and the nitrogen cycle (Ulyshen 2015). In addition, saproxylic and leaf-dwelling beetles can be important prey for insectivorous vertebrates (e.g. Tillon et al. 2016; Koenig and Liebhold 2017), and changes in beetle community composition may therefore have cascading effects on the food web (e.g. Koenig and Liebhold 2017). From a conservation standpoint, the increase in species richness for the xylophagous and phyllophagous beetle communities suggests that declining stands might enhance forest biodiversity. Decline especially promoted saproxylic species. This community is particularly sensitive to the intensification of management practices involving the extraction of weakened or decaying wood material, and consequently includes several rare and protected species (Grove 2002; Seibold et al. 2015). The accumulation of suitable habitats and resources for this community in declining stands may then counterbalance the adverse effects of intensive management. The increase in abundance and/or biomass of xylophagous and phyllophagous beetles also resulted in an overall increase in beetle abundance and biomass in the declining stands. This also suggests that forest decline may mitigate the reduction in insect biomass recently reported in European forests, in intensively managed landscapes (Seibold et al. 2019), at least if the increase in resources and structural complexity persists over time (Winter et al. 2015). In this regard, increases in species richness, abundance and biomass of xylophagous species at the stand scale, as in our study, might prove to be ephemeral (Winter et al. 2015).

In our forests, in the CAP sampling year explained a greater percentage of inertia than did decline level for all guilds considered. This strong year effect could result from high inter-annual variations in beetle abundance and/or occurrence, but may also incorporate multiple methodological factors (i.e., (i) slight variations in sampling periods, (ii) changes in monitored plots and (iii) modifications in the protocol of decline characterization at the plot level). Marked variations in beetle abundance and biomass occurred on plots and at periods that were consistently monitored throughout the three years of survey, and between years (i.e., 2016 and 2017) when the protocol of decline characterization was identical (data not shown). This rather supports the hypothesis that the year effect mainly results from marked inter-annual variations in beetle abundance and community composition. Such fluctuations in population and community abundances are commonly observed in temperate forests (e.g. Stange et al. 2011). A longer monitoring period on the same plots would be necessary to identify the factors contributing to the between-year variations we observed.

Green Lindgren traps, placed at the canopy level, have proven to be effective in collecting leaf-dwelling beetles. These traps were specifically designed to collect *Agrilus planipennis* Fairmaire (Francese et al. 2011), but have also allowed researchers to collect North American and European Agrilinae species (Petrice and Haack 2015; Rassati et al. 2019). During our survey, all the Agrilinae species associated with oaks in France (i.e. *Agrilus* sp., *Coraebus* sp. and *Meliboeus* sp.) were captured, except for *Agrilus grandiceps hemiphanes* Marseul, a rare Mediterranean species, and *Coraebus florentinus* Herbst. The latter species had previously been collected in the Vierzon forest, and typical shoot browning resulting from its larval activity has already been reported there. The species might have been present but at too low population density for detection, or it might not have been attracted by our traps. We also collected quite diverse communities of phyllophagous and seminiphagous weevil species in our green Lindgren traps, in large amounts for some species. These species were significantly more attracted to green traps than to purple ones, which is congruent with the attraction to green substrates reported for other phytophagous weevils (e.g. Cross et al. 1976; Gadi and Reddy 2014). Overall, this suggests that green Lindgren traps are attractive to phyllobiont species in general, and confirms the tool’s utility when investigating canopy-dwelling beetles associated with foliage.

## Conclusion

Our three-year survey in a declining forest allowed us to detect significant effects of decline on different canopy-dwelling species and guilds, in spite of strong inter-annual variations and a limited spatial extent, the survey being performed in two adjacent forests. Overall, decline had a positive effect on the abundance and biomass of beetles, but contrasted variations were observed at the species or guild levels, with positive effects for saproxylic and generalist phyllophagous species, null effects for seminiphagous species and negative effects for specialist phyllophagous species. These results call for studies conducted at larger spatial and temporal scales to assess the functional outcomes of the unprecedented level of forest decline expected to affect Europe, and to propose management strategies for conservation biologists.

## Contributions of the co-authors

Conceptualization: A. Sallé & C. Bouget; Methodology: G. Parmain, B. Nusillard, X. Pineau; Data acquisition: G. Parmain, B. Nusillard, X. Pineau, R. Brousse, T. Fontaine-Guenel, R. Ledet, C. Vincent-Barbaroux; Writing – original draft: A. Sallé & C. Bouget; Writing – review & editing: A. Sallé, C. Bouget, G. Parmain, C. Vincent-Barbaroux; Supervision: A. Sallé, Funding acquisition: A. Sallé

## Acknowledgements

We thank C. Moliard (INRAE) for his technical assistance. We are grateful to O. Rose (Ciidae), T. Noblecourt (Scolytinae), F. Soldati (Tenebrionidae, Carabidae), T. Barnouin (Elateridae (pars), Ptinidae (pars)), O. Courtin (Scraptiidae, Mordellidae), Y. Gomy (Histeridae), and C. Sallé (Curculioninae) for their help with the identifications. We are also grateful to the National Forestry Office (Office National des Forêts) and the Forest Health Service (Département de la Santé des Forêts), with special thanks to A. Hachette, F.-X. Saintonge and D. Baudet for their field assistance.

## Data availability

The datasets generated during and/or analyzed during the current study are not publicly available due to further analyses on the data but are available from the corresponding author on reasonable request.

---

**Figure S1.**
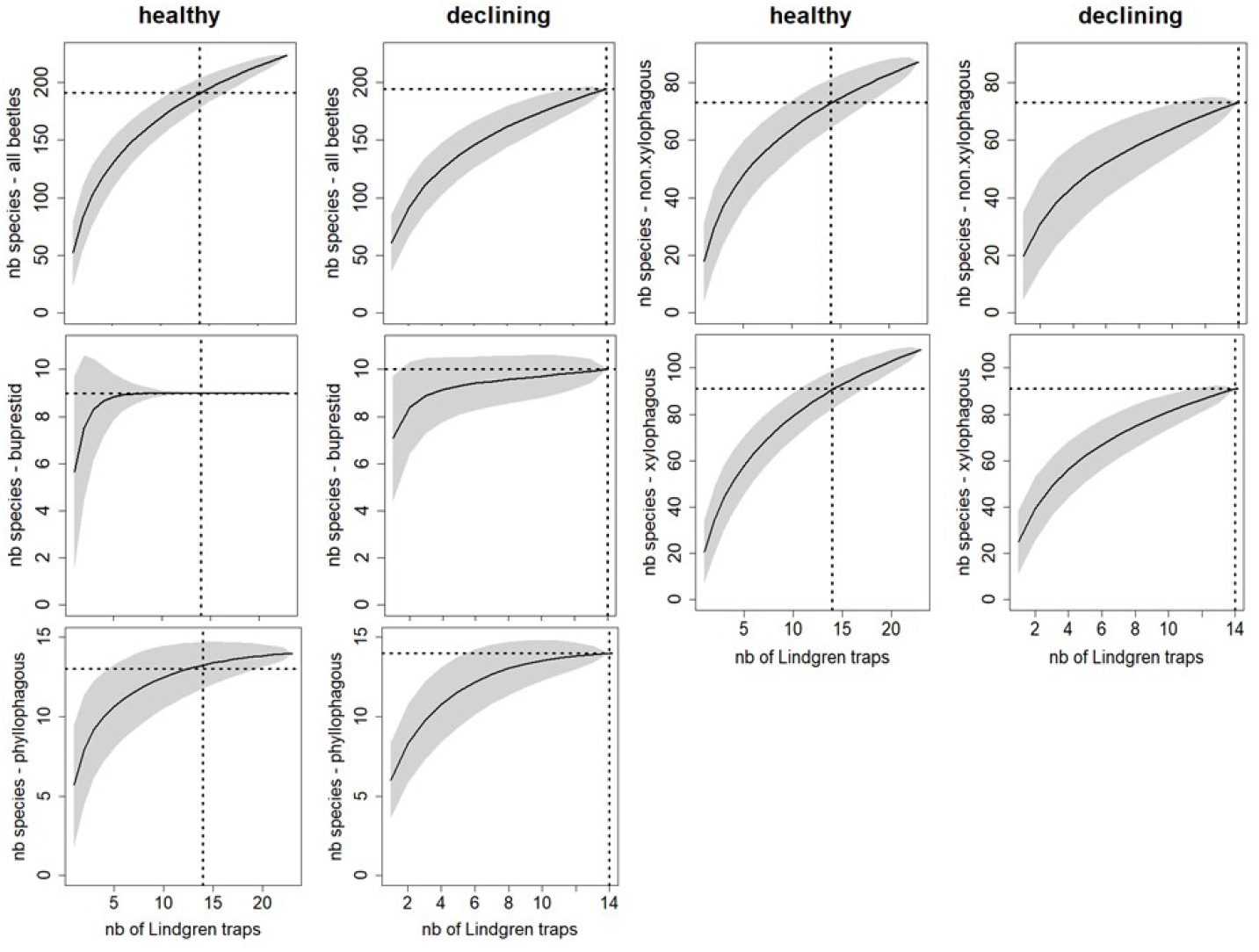
Cumulative interpolated rarefaction of species richness with sample size (sampling without replacement) by dieback level at tree scale. Vertical dashed line = standard interpolated sample size, horizontal dashed line = species richness estimate at the standardized sample size, and grey area = standard deviation of species richness estimate.

**Table S1:**
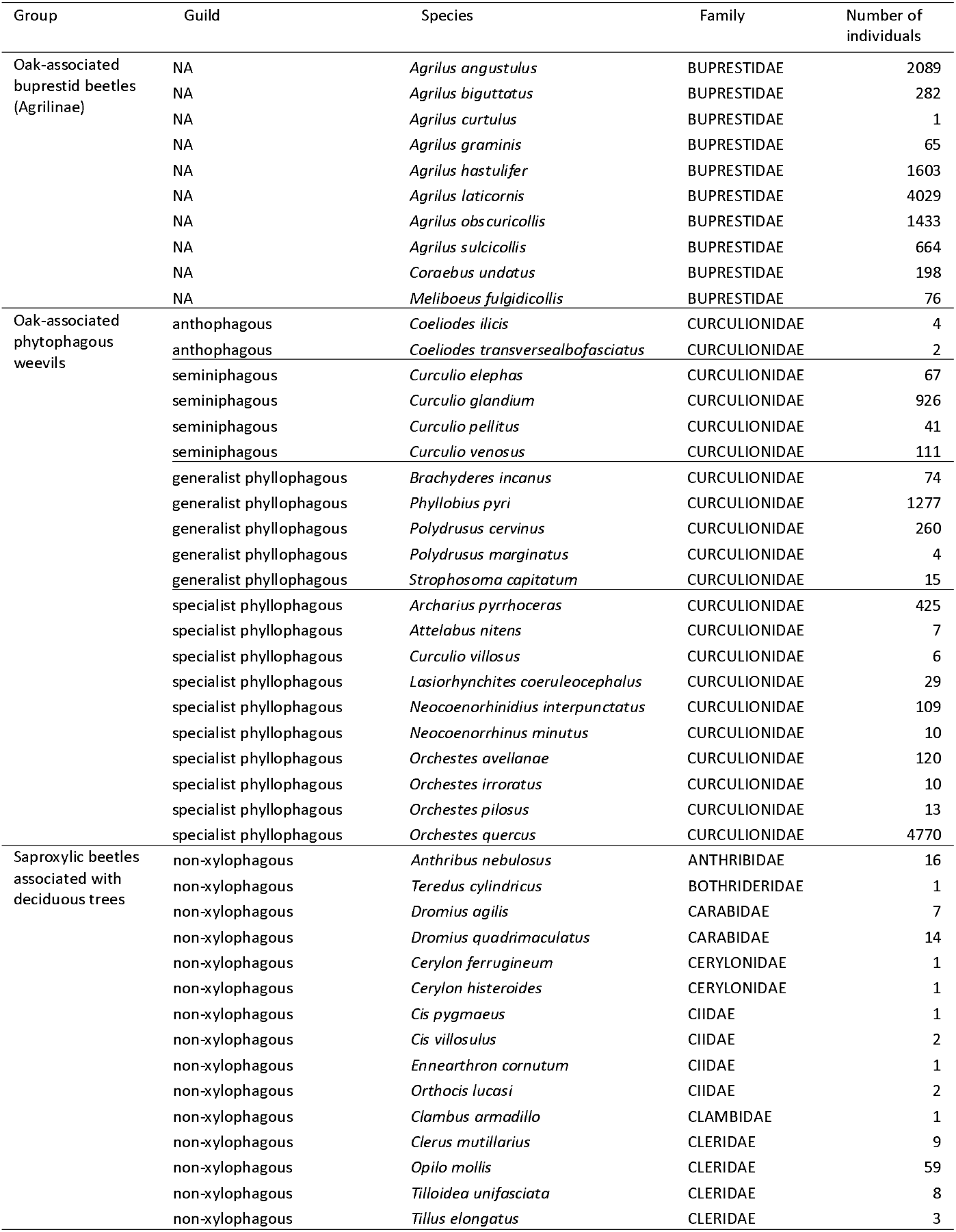
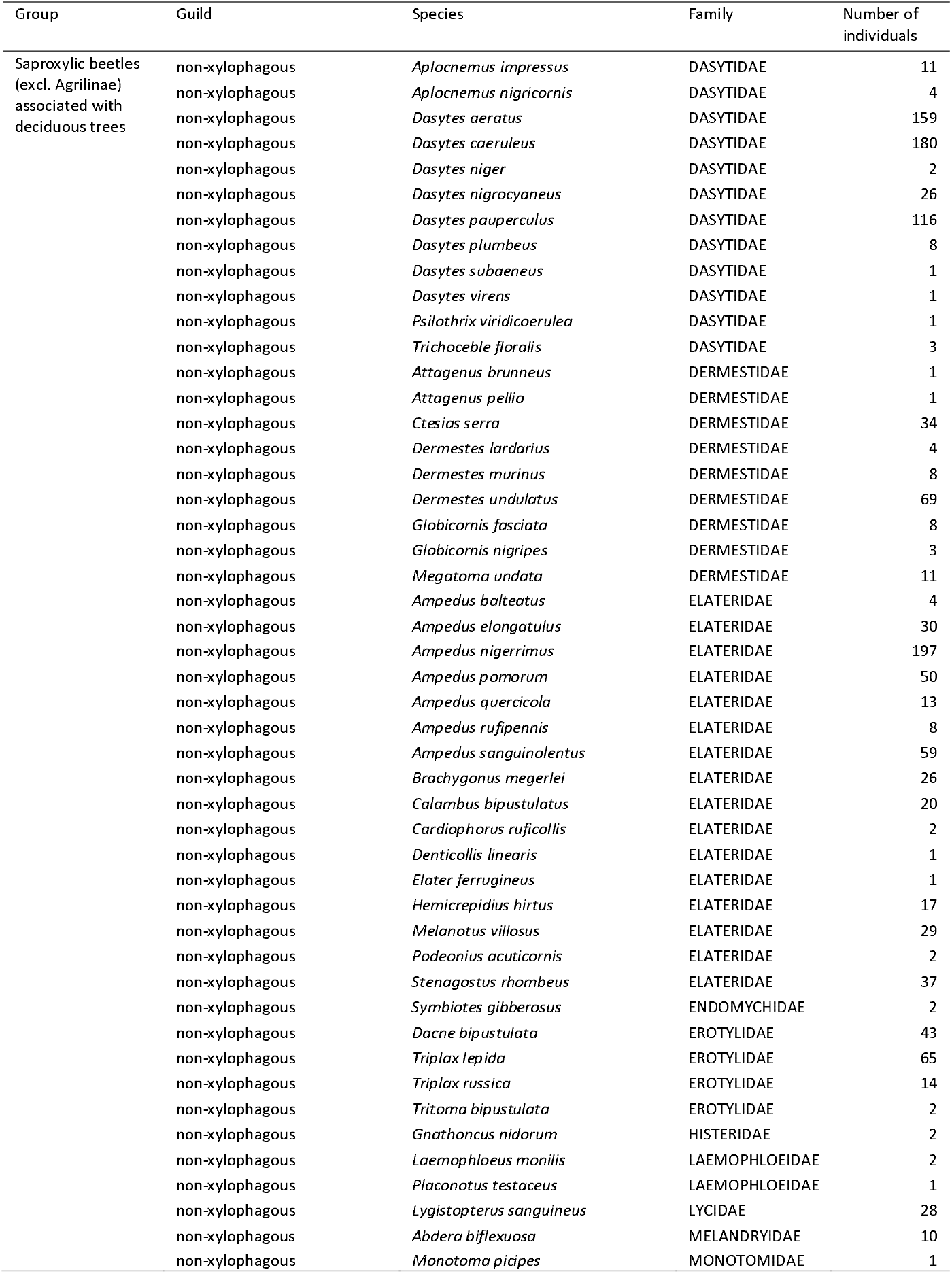
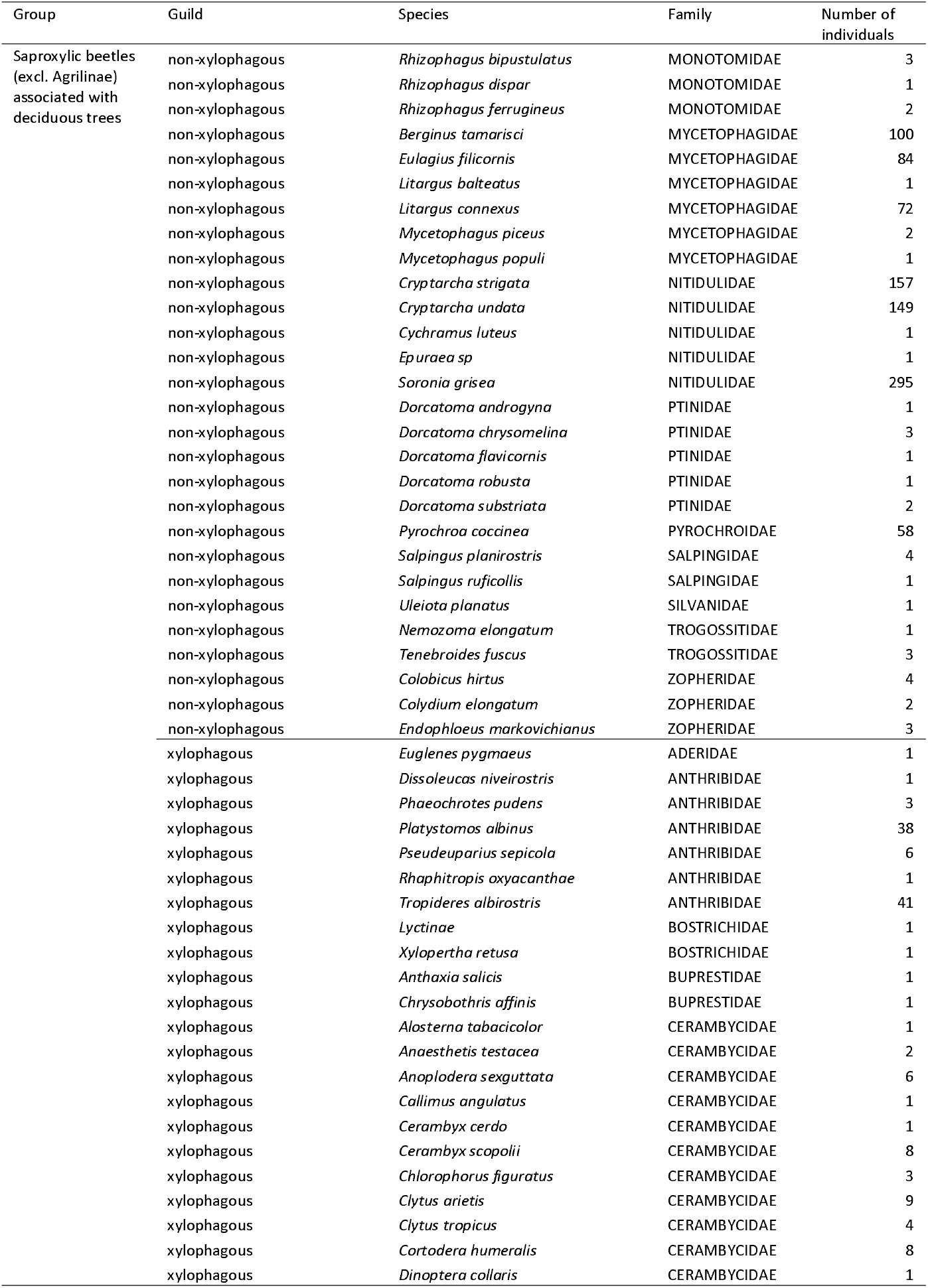
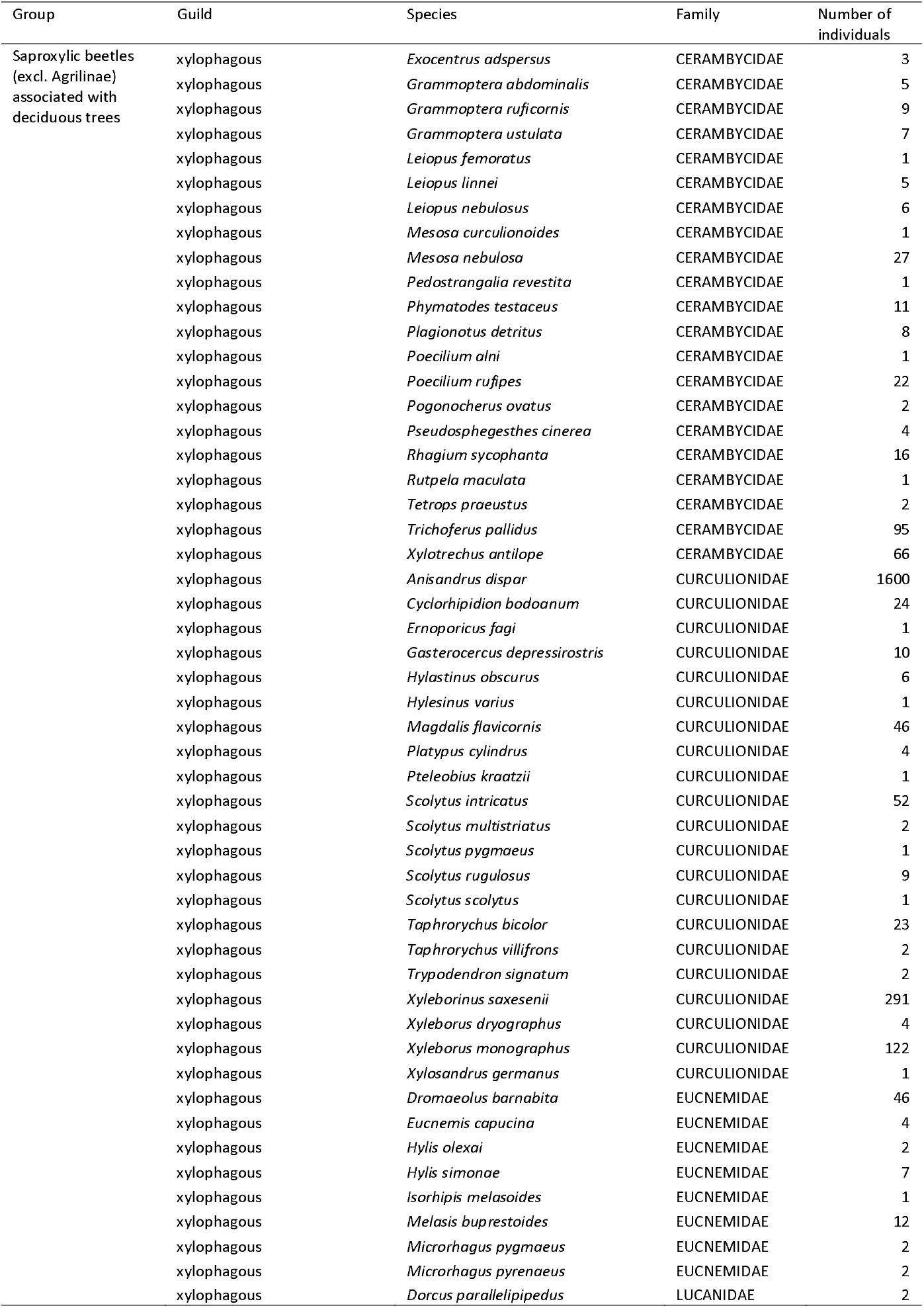
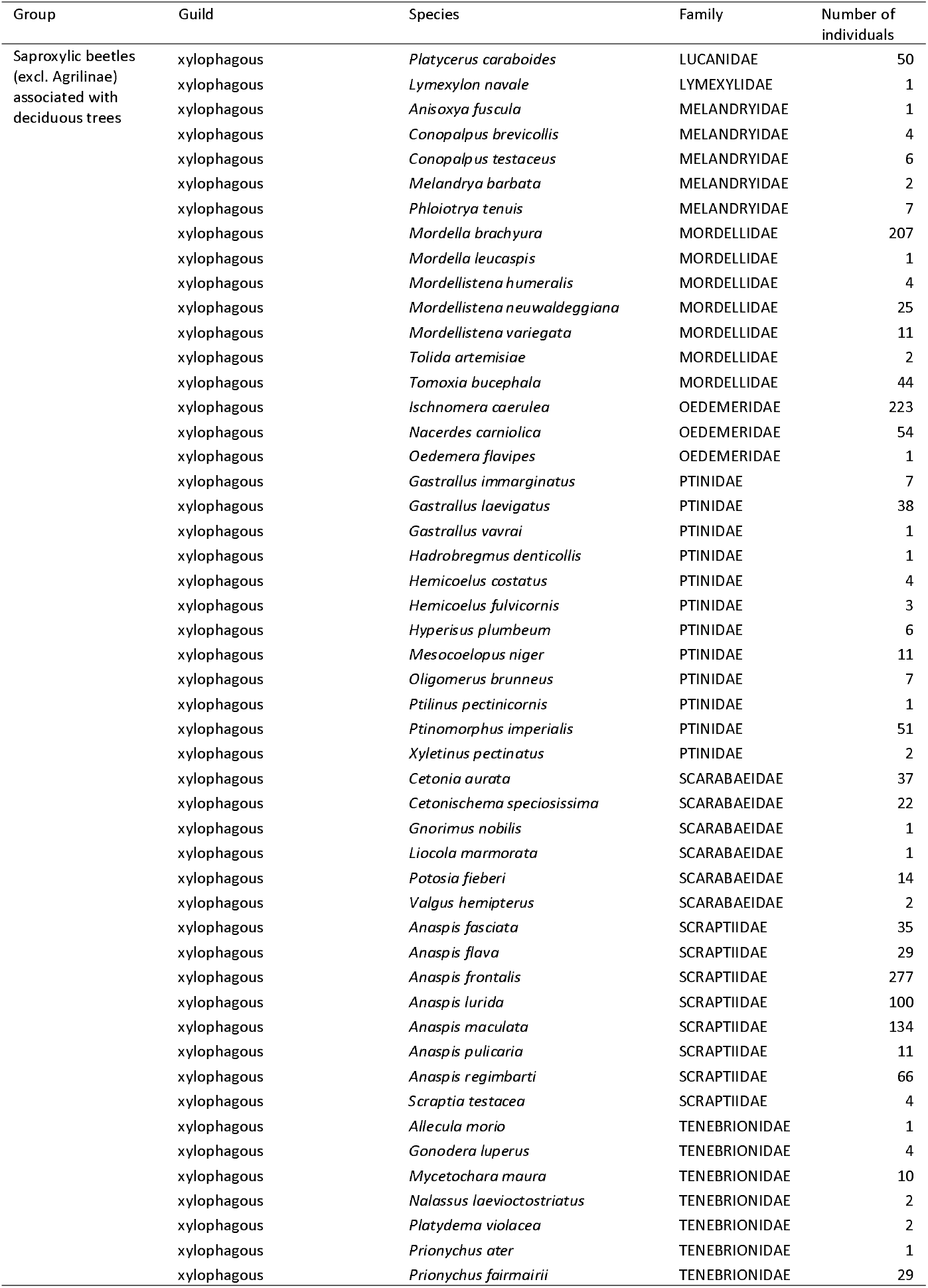
list of all the beetle species collected and used for the data analyses with their group, their guild and their abundance

## References

Allen C D, Macalady A K, Chenchouni H, Bachelet D, McDowell N, Vennetier M, Kitzberger T, Rigling A, Breshears D D, Hogg E H, Gonzalez P, Fensham R, Zhang Z, Castro J, Demidova, N, Lim J-H, Allard G, Running S W, Semerci A, Cobb N (2010) A global overview of drought and heat-induced tree mortality reveals emerging climate change risks for forests. For Ecol Manage 259: 660–684.

Anderson M J, Willis T J (2003) Canonical analysis of principal coordinates: a useful method of constrained ordination for ecology. Ecol 84: 511–525.

Bouget C, Brustel H, Noblecourt T, Zagatti P (2019) Les Coléoptères saproxyliques de France: Catalogue écologique illustré. Muséum National d’Histoire Naturelle, Paris, 744p.

Bouget C, Brin A, Brustel H (2011) Exploring the “last biotic frontier”: are temperate forest canopies special for saproxylic beetles? For Ecol Manage 261: 211–220.

Brin A, Bouget C, Brustel H, Jactel H (2011) Diameter of downed woody debris does matter for saproxylic beetle assemblages in temperate oak and pine forests. J Insect Conserv 15: 653–669.

Brown N, Jeger M, Kirk S, Williams D, Xu X, Pautasso M, Denman S (2017) Acute Oak Decline and *Agrilus biguttatus:* the co-occurrence of stem bleeding and D-shaped emergence holes in Great Britain. Forests, 8: 87.

Brück-Dyckhoff C, Petercord R, Schopf R (2019) Vitality loss of European beech (Fagus sylvatica L.) and infestation by the European beech splendour beetle *(Agrilus viridis* L., Buprestidae, Coleoptera). Forest Ecology and Management, 432: 150–156.

Carnicer J, Coll M, Ninyerola M, Pons X, Sanchez G, Penuelas J (2011) Widespread crown condition decline, food web disruption, and amplified tree mortality with increased climate change-type drought. Proc Nat Acad Sci 108: 1474–1478.

Cross W H, Mitchell H C, Hardee D D (1976) Boll weevils: Response to light sources and colors on traps. Environ Entomol 5: 565–57.

De Frenne P, Rodríguez-Sánchez F, Coomes D A, Baeten L, Verstraeten G, Vellend M, Bernhardt-Römermann M, Brown C D, Brunet J, Cornelis J, Decocq G M, Dierschke H, Eriksson O, Gilliam F S, Hédl R, Heinken T, Hermy M, Hommel P, Jenkins M A, Kelly D L, Kirby K J, Mitchell F J G, Naaf T, Newman M, Peterken G, Petrík P, Schultz J, Sonnier G, Van Calster H, Waller D M, Walther G-R, White P S, Woods K D, Wulf M, Graae B J, Verheyen K (2013) Microclimate moderates plant responses to macroclimate warming. Proc. Nat. Acad. Sci., 110: 18561–18565.

De Frenne P, Zellweger F, Rodriguez-Sanchez F, Scheffers B R, Hylander K, Luoto M, Vellend M, Verheyen K, Lenoir J (2019) Global buffering of temperatures under forest canopies. Nat Ecol Evol 3: 744–749.

Delatour C (1983) Les dépérissements de chênes en Europe. Rev For Fra 4: 265–282.

Denman S, Brown N, Kirk S, Jeger M, Webber J (2014) A description of the symptoms of Acute Oak Decline in Britain and a comparative review on causes of similar disorders on oak in Europe. Forestry 87: 535–551.

Douzon G (2006) Le point sur les dépérissements des chênes pédonculés en forêt de Vierzon. Département de la santé des forêts, Bilan de la santé des forêts en 2005, 4 p.

Dufrêne M, Legendre P (1997) Species assemblages and indicator species: the need for a flexible asymmetrical approach. Ecol Monogr 67: 345–366.

Forkner R E, Marquis R J, Lill J T (2004) Feeny revisited: condensed tannins as anti-herbivore defences in leaf-chewing herbivore communities of *Quercus*. Ecol Entomol 29: 174–187.

Francese J A, Fraser I, Lance D R, Mastro V C (2011) Efficacy of multifunnel traps for capturing emerald ash borer (Coleoptera: Buprestidae): effect of color, glue, and other trap coatings. J Econ Entomol 104: 901–908.

Gadi N, Reddy G V (2014) Are sweetpotato weevils (Coleoptera: Brentidae) differentially attracted to certain colors? Ann Entomol Soc Am 107: 274–278.

Ganihar S R (1997) Biomass estimates of terrestrial arthropods based on body length. J Biosci 22: 219–224.

Gely C, Laurance S G, Stork N E (2019) How do herbivorous insects respond to drought stress in trees? Biol Rev doi: 10.1111/brv.12571.

Grove S J (2002) Saproxylic insect ecology and the sustainable management of forests. Ann Rev Ecol Syst 33: 1–23.

Günthardt-Goerg M S, Kuster T M, Arend M, Vollenweider P (2013) Foliage response of young central European oaks to air warming, drought and soil type. Plant Biol 15: 185–197.

Gutbrodt B, Mody K, Dorn S (2011) Drought changes plant chemistry and causes contrasting responses in lepidopteran herbivores. Oikos 120: 1732–1740.

Haack R A, Benjamin D M (1982) The biology and ecology of the twolined chestnut borer, *Agrilus bilineatus* (Coleoptera: Buprestidae), on oaks, *Quercus* spp., in Wisconsin. Can Entomol 114: 385–396.

Hardwick S R, Toumi R, Pfeifer M, Turner E C, Nilus R, Ewers R M (2015) The relationship between leaf area index and microclimate in tropical forest and oil palm plantation: Forest disturbance drives changes in microclimate. Agric For Meteorol 201: 187–195.

Heitzman E, Grell A, Spetich M, Starkey D (2007) Changes in forest structure associated with oak decline in severely impacted areas of northern Arkansas. South J Appl For 31: 17–22.

Herms D A, Mattson W J (1992) The dilemma of plants: to grow or defend. Q Rev Biol 67: 283–335.

Houston D R (1981) Stress triggered tree diseases: the diebacks and declines. USDA For Serv Rep INF-41-81.

Hu B, Simon J, Rennenberg H (2013) Drought and air warming affect the species-specific levels of stress-related foliar metabolites of three oak species on acidic and calcareous soil. Tree Physiol 33: 489–504.

IPCC (2013) IPCC, 2013: summary for policymakers. In: Stocker TF Qin D Plattner G-K et al. (eds) Climate change 2013: the physical science basis. Cambridge University Press, Cambridge (UK) and New York (USA).

Ishii H T, Tanabe S I, Hiura T (2004) Exploring the relationships among canopy structure, stand productivity, and biodiversity of temperate forest ecosystems. For Sci 50: 342–355.

Jennings D E, Taylor P B, Duan J J (2014) The mating and oviposition behavior of the invasive emerald ash borer *(Agrilus planipennis)*, with reference to the influence of host tree condition. J Pest Sci 87: 71–78.

Kaplan I, Denno R F (2007) Interspecific interactions in phytophagous insects revisited: a quantitative assessment of competition theory. Ecol Lett 10: 977–994.

Koenig W D, Liebhold A M (2017) A decade of emerald ash borer effects on regional woodpecker and nuthatch populations. Biol Invasions, 19: 2029–2037.

Kozák D, Svitok M, Wiezik M, Mikoláš M, Thorn S, Buechling A, Hofmeister J, Matula R, Trotsiuk V, Bace V, Begovic K, Cada V, Dusatko M, Frankovic M, Horak J, Janda P, Kameniar O, Nagel T A, Pettit J L, Pettit J M, Synek M, Wiezikova A, Svoboda M (2020) Historical disturbances determine current taxonomic, functional and phylogenetic diversity of saproxylic beetle communities in temperate primary forests. Ecosystems, doi 10.1007/s10021-020-00502-x.

Lämke J S, Unsicker S B (2018) Phytochemical variation in treetops: causes and consequences for tree-insect herbivore interactions. Oecologia, 187: 377–388.

Larsson S (1989) Stressful times for the plant stress: insect performance hypothesis. Oikos 56: 277–283.

Liebhold A M, Brockerhoff E G, Kalisz S, Nuñez M A, Wardle D A, Wingfield M J (2017) Biological invasions in forest ecosystems. Biol Invasions 19: 3437–3458.

Manion P D (1981) Tree disease concepts. Prentice-Hall, Englewood Cliffs.

Marçais B, Desprez-Loustau M L (2014) European oak powdery mildew: impact on trees, effects of environmental factors, and potential effects of climate change. Ann For Sci 71: 633–642.

Martel J, Mauffette Y (1997) Lepidopteran communities in temperate deciduous forests affected by forest decline. Oikos 78: 48–56.

Millar C I, Stephenson N L (2015) Temperate forest health in an era of emerging megadisturbance. Sci 349: 823–826.

Montgomery M E, Wargo P M (1983) Ethanol and other host-derived volatiles as attractants to beetles that bore into hardwoods. J Chem Ecol, 9: 181–190.

Moraal L G, Hilszczanski J (2000) The oak buprestid beetle, *Agrilus biguttatus* (F.) (Col., Buprestidae), a recent factor in oak decline in Europe. J Pest Sci 73: 134–138.

Nageleisen L M (2005). Dépérissement du hêtre: présentation d’une méthode symptomatologique de suivi. Rev For Fr 57: 255–262.

Oszako T (2000) Oak declines in Europe’s forest – history, causes and hypothesis, in Oszako T, Delatour C (Eds.), Recent advances on oak health in Europe. Forest Research Institute, Warsaw, Poland, pp. 11–40.

Petrice T R, Haack R A (2014) Biology of the European oak borer in Michigan, United States of America, with comparisons to the native twolined chestnut borer. Can Entomol 146: 36–51.

Petrice T R, Haack R A (2015) Comparison of different trap colors and types for capturing adult *Agrilus* (Coleoptera: Buprestidae) and other buprestids. Great Lakes Entomol 48: 45–66.

Pincebourde S, Sinoquet H, Combes D, Casas J (2007) Regional climate modulates the canopy mosaic of favourable and risky microclimates for insects. J Anim Ecol 76: 424–438.

Plewa R, Jaworski T, Hilszczański J, Horák J (2017) Investigating the biodiversity of the forest strata: The importance of vertical stratification to the activity and development of saproxylic beetles in managed temperate deciduous forests. For Ecol Manage 402: 186–193.

Poole E M, Ulyshen M D, Horn S, Cram M, Olatinwo R, Fraedrich S (2019) Biology and distribution of *Agrilus macer* LeConte (Coleoptera: Buprestidae), a species associated with sugarberry *(Celtis laevigata* Willd.) mortality in the southeastern USA. Ann For Sci 76: 7.

R Core Team (2018) R: A language and environment for statistical computing. R Foundation for Statistical Computing, Vienna, Austria. URL http://www.R-project.org/

Rassati D, Marini L, Marchioro M, Rapuzzi P, Magnani G, Poloni R, Di Giovanni F, Mayo P, Sweeney J (2019) Developing trapping protocols for wood-boring beetles associated with broadleaf trees. J Pest Sci 92: 267–279.

Redilla K M, McCullough D G (2017) Species assemblage of buprestid beetles in four hardwood cover types in Michigan. Can J For Res 47: 1131–1139.

Saintonge F-X, Nageleisen L-M (2001) Le rôle des agriles dans le dépérissement des chênes: observations récentes en Alsace. In: ONF, FVA (Eds.), Insectes corticoles et dépérissement des chênes, pp. 33–42.

Sallé A, Nageleisen L-M, Lieutier F (2014) Bark and wood boring insects involved in oak declines in Europe: current knowledge and future prospects in a context of climate change. For Ecol Manage 328: 79–93.

Sánchez-Osorio I, López-Pantoja G, Tapias R, Pareja-Sánchez E, Domínguez L (2019) Monoterpene emission of *Quercus suber* L. highly infested by *Cerambyx welensii* Küster. Ann For Sci 76: 98.

Seibold S, Brandl R, Buse J, Hothorn T, Schmidl J, Thorn S, Müller J (2015) Association of extinction risk of saproxylic beetles with ecological degradation of forests in Europe. Conserv Biol, 29: 382–390.

Seibold S, Gossner M M, Simons N K, Blüthgen N, Müller J, Ambarli D, Ammer C, Bauhus J, Fischer M, Habel J C, Linsenmair K E, Nauss T, Penone C, Prati D, Schall P, Schulze E-D, Vogt J, Wöllauer S, Weisser W W (2019) Arthropod decline in grasslands and forests is associated with landscape-level drivers. Nat 574: 671–674.

Seidl R, Thom D, Kautz M, Martin-Benito D, Peltoniemi M, Vacchiano G, Wild J, Ascoli D, Petr M, Honkaniemi J, Lexer M J, Trotsiuk V, Mairota P, Svoboda M, Fabrika M, Nagel T A, Reyer C P O (2017) Forest disturbances under climate change. Nat Clim Change 7: 395–402.

Sinclair, W.A., 1967. Decline of hardwoods: possible causes, in: International shade tree conference, pp. 17–32.

Sonesson K, Drobyshev I (2010) Recent advances on oak decline in southern Sweden. Ecol Bull 53: 197–208.

Southwood T R E (1961) The number of species of insect associated with various trees. J Anim Ecol 30: 1–8.

Spetich M A (2007) Down deadwood dynamics on a severely impacted oak decline site. USDA For Serv Gen Tech Rep SRS-101: 206–213.

Stange E E, Ayres M P, Bess J A (2011) Concordant population dynamics of Lepidoptera herbivores in a forest ecosystem. Ecography 34: 772–779.

Stokland J N, Siitonen J, Jonsson B G (2012) Biodiversity in Dead Wood. Cambridge University press.

Stork N E, Grimbacher P S (2006) Beetle assemblages from an Australian tropical rainforest show that the canopy and the ground strata contribute equally to biodiversity. Proc Roy Soc B: Biol Sci 273: 1969–1975.

Thomas F M, Blank R, Hartmann G (2002) Abiotic and biotic factors and their interactions as causes of oak decline in Central Europe. For Pathol 32: 277–307.

Tiberi R, Branco M, Bracalini M, Croci F, Panzavolta T (2016) Cork oak pests: a review of insect damage and management. Ann For Sci 73: 219–232.

Tillon L, Bouget C, Paillet Y, Aulagnier S (2016) How does deadwood structure temperate forest bat assemblages? Eur J For Res 135: 433–449.

Tschanz B, Schmid E, Bacher S (2005) Host plant exposure determines larval vulnerability-do prey females know? Funct Ecol 19: 391–395.

Ulyshen M D (2015) Insect-mediated nitrogen dynamics in decomposing wood. Ecol Entomol 40: 97–112.

Ulyshen M D (2011) Arthropod vertical stratification in temperate deciduous forests: implications for conservation-oriented management. For Ecol Manage 261: 1479–1489.

Vodka Š, Cizek L (2013) The effects of edge-interior and understorey-canopy gradients on the distribution of saproxylic beetles in a temperate lowland forest. For Ecol Manage 304: 33–41.

Winter M B, Ammer C, Baier R, Donato D C, Seibold S, Müller J (2015) Multi-taxon alpha diversity following bark beetle disturbance: evaluating multi-decade persistence of a diverse early-seral phase. For Ecol Manage 338: 32–45.

Yamasaki M, Kikuzawa K (2003) Temporal and spatial variations in leaf herbivory within a canopy of *Fagus crenata*. Oecologia 137: 226–232.

